# Neurexin1α differentially regulates synaptic efficacy within striatal circuits

**DOI:** 10.1101/2020.05.31.125575

**Authors:** M. Felicia Davatolhagh, Marc V. Fuccillo

**Affiliations:** Neuroscience Graduate Group, University of Pennsylvania, Philadelphia, PA 19104, USA; Department of Neuroscience, Perelman School of Medicine, University of Pennsylvania, Philadelphia, PA 19104, USA

**Keywords:** corticostriatal, thalamostriatal, synaptic transmission, Neurexin1α, prefrontal cortex, striatum

## Abstract

Mutations in genes essential for shared aspects of synaptic function throughout the CNS, such as the presynaptic adhesion molecule Neurexin1α (Nrxn1α), are strongly implicated in neuropsychiatric pathophysiology. As the input nucleus of the basal ganglia, the striatum integrates diverse excitatory projections governing cognitive and motor control, and its functional impairment underlies neuropsychiatric disorders. While prior work has emphasized Neurexins’ contributions to synaptic transmission in hippocampus and brainstem, their function in striatal circuits remains unstudied. As Nrxn1α is highly expressed at striatal inputs, we employed optogenetic-mediated afferent recruitment of dorsal prefrontal cortex-dorsomedial striatal (DMS) connections, uncovering a decrease in net synaptic strength specifically onto indirect pathway spiny neurons in both Nrxn1α^+/-^ and Nrxn1α^-/-^ mice, driven by reductions in transmitter release probability. In contrast, thalamic excitatory inputs to DMS demonstrated relatively normal excitatory synaptic strength. These findings suggest that dysregulation of Nrxn1α modulates striatal function in an input and target-specific manner.

## Introduction

The striatum, as the input nucleus of the basal ganglia, integrates diverse neuronal projections governing cognitive and motor control (Balleine et al., 2009; Graybiel et al., 1994; Packard and Knowlton, 2002), and serves behavioral functions whose impairment is linked to neuropsychiatric disorders (Fuccillo, 2016; Yin and Knowlton, 2006). The principal striatal cell type, spiny projection neurons (SPNs), which is distinguished by expression of D1 or D2 dopamine receptors (Gerfen et al., 1990), receives widespread forebrain glutamatergic input from cortex and thalamus (Ding et al., 2008; Pan et al., 2010). These excitatory projections have topographical organization (Deng et al., 2015; Gerfen, 1989; Hunnicutt et al., 2016), extensive molecular diversity and distinct synaptic properties (Ding et al., 2008; Smeal et al., 2008). This complexity has been a persistent obstacle in understanding how molecular dysfunction contributes to disease-relevant striatal circuit abnormalities.

As the number of neuropsychiatric disease-associated mutations grows, candidate genes have been parsed according to putative functions (Chang et al., 2015; Schizophrenia Working Group of the Psychiatric Genomics, 2014) and spatio-temporal cellular expression patterns (Willsey et al., 2013). Functionally, there is an overrepresentation of genes encoding DNA regulatory proteins, neuronal signaling regulators and molecules essential for development and function of synapses, including synaptic adhesion proteins (Sudhof, 2008, 2018; Willsey and State, 2015). Spatially, cortico-striato-thalamic (CST) circuits reliably exhibit enrichment in expression of candidate genes for both autism spectrum disorder (ASD) and schizophrenia (Chang et al., 2015; Willsey et al., 2013).

Thus far, analysis of synaptic adhesion molecule function within CST circuits has centered on Neuroligin (NL) family postsynaptic proteins. Dorsal striatal recordings of NL1 knock-out (KO) mice revealed a direct pathway dSPN-specific reduction in the ratio of N-methyl-D-aspartate receptor (NMDAR) to α-amino-3-hydroxy-5-methyl-4-isoxazolepropionic acid receptor (AMPAR)-mediated currents, which likely contributed to increased grooming in these KOs (Blundell et al., 2010). In the nucleus accumbens, NL3 disruption selectively impaired synaptic inhibition of dSPNs, contributing to enhanced motor learning (Rothwell et al., 2014). Despite their extensive disease association and complex behavioral picture, the function of Neurexins (Nrxns), Neuroligin’s presynaptic partners, within striatal circuits remains unexplored.

In mammals, Nrxns are encoded by three genes under control of alpha (α) and beta (β) promoters. While they are key molecules mediating synapse organization and calcium-triggered neurotransmitter release (Missler et al., 2003), specific functions depend on Nrxn isoform, brain region and synapse subtype. In hippocampal subiculum, Nrxn3 functions trans-synaptically to maintain postsynaptic AMPAR content, while β-Nrxns regulate release probability through modulation of endocannabinoid signaling (Anderson et al., 2015). Within the Nrxn family, Nrxn1α exhibits a disproportional neuropsychiatric disease association, with numerous heterozygous loss-of-function alleles found in ASD, schizophrenia, Tourette syndrome, and obsessive compulsive disorder (OCD) (Bucan et al., 2009; Ching et al., 2010; Dabell et al., 2013; Kim et al., 2008; Kirov et al., 2009; Reichelt et al., 2012). Despite these findings, Nrxn1α mutations have only been examined in hippocampus and analysis of heterozygotes has been overlooked (Etherton et al., 2009).

Here, we assessed the functional integrity of striatal circuits in Nrxn1α^+/-^ and Nrxn1α^-/-^ mice using optogenetic tools to specifically interrogate excitatory afferents into striatum. Using a combined field/whole-cell recording configuration we found a decrease in dPFC-DMS synaptic efficacy specifically onto the indirect iSPN pathway. This reduction in excitatory strength resulted from a reduction in glutamate release probability and decreased dPFC-iSPN synaptic drive across a broad range of naturalistic input frequencies. In contrast, another key excitatory striatal circuit, parafascicular thalamic (PFas) inputs to DMS, exhibited normal presynaptic function but altered NMDAR content onto both SPN subtypes. Taken together, we identify input and target-specific alterations in striatal circuits of mice with both heterozygous and homozygous mutations in Nrxn1α.

## Results

### Neurexin1α Deletion alters Spontaneous Excitatory Synaptic Transmission within Dorsal Striatum

To assess whether Neurexin1α deletion impacts excitatory synaptic function onto dorsal striatal SPNs, we performed whole-cell voltage-clamp recordings in the dorsomedial striatum (DMS) of acute slices taken from Neurexin1α heterozygote and homozygous knock-out (KO) animals. Mice were crossed onto the Drd1a-tdTomato BAC transgenic line, permitting identification of dSPNs and putative iSPNs (Ade et al., 2011; Choi et al., 2019). We assessed global excitatory synaptic function by recording TTX-insensitive spontaneous miniature excitatory postsynaptic currents (mESPCs) (Fig1A,C). For dSPNs, we noted no genotypic differences in mEPSC amplitude (WT: 18.72 ± 0.7 pA; n = 23/7, Het: 18.74 ± 0.7 pA; n = 18/6, KO: 19.44 ± 0.6 pA; n = 24/7, ANOVA), but an increase in mEPSC frequency (WT: 2.94 ± 0.3 Hz, Het: 3.71 ± 0.3 Hz, KO: 4.97 ± 0.3 Hz, Dunnett’s test) in Nrxn1α-KO mice, with an intermediate increase in Nrxn1α-Hets (Fig.1A). In contrast, no changes in the amplitude (WT: 19.32 ± 0.8 pA; n = 20/9, Het: 19.59 ± 1.1 pA; n = 15/6, KO: 18.25 ± 0.5 pA; n = 22/7, 1-way ANOVA) or frequency of mEPSCs were observed for iSPNs (WT: 2.863 ± 0.3 Hz, Het: 3.89 ± 0.3 Hz, KO: 3.43 ± 0.3 Hz, ANOVA) (Fig.1C). To broadly assess spontaneous inhibitory transmission, we also recorded miniature inhibitory postsynaptic currents (mIPSCs). We did not observe statistically significant differences for either mIPSC frequency or amplitude across all tested genotypes, regardless of the subtype of recorded SPNs (Supp.Fig.1A,B).

**Figure 1.**
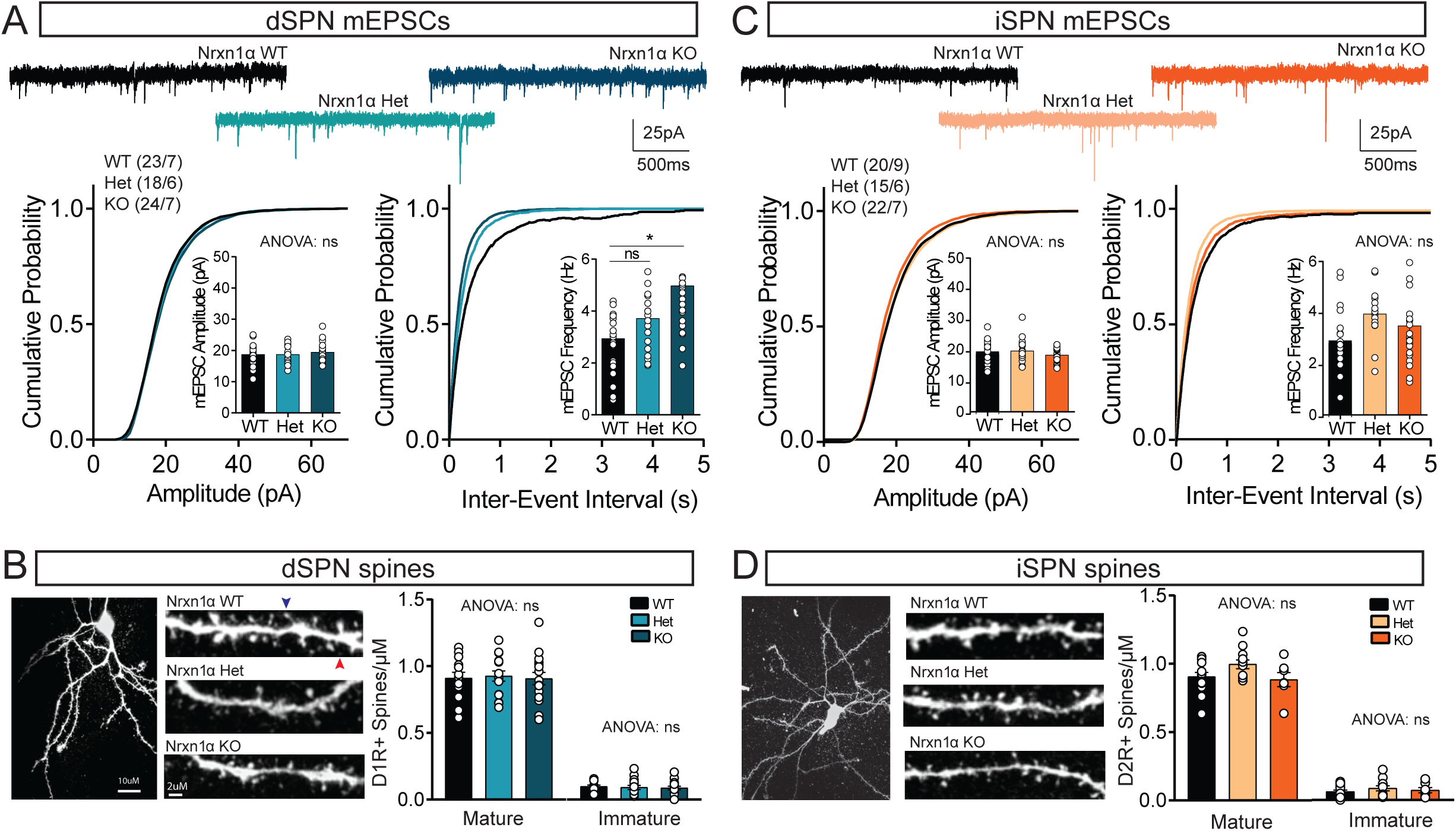
Nrxn1α heterozygous and homozygous animals display altered spontaneous synaptic transmission without a change in overall number of excitatory synapses. (A and C) Representative traces of mEPSCs WT, Nrxn1α Het, and Nrxn1α KO cells (top), cumulative distribution of mEPSC amplitude (lower left; inset shows average mEPSC amplitude), and cumulative distribution of inter-event intervals (lower right; inset shows average mEPSC frequency) in WT (n = 23; N = 7), Nrxn1α Het (n = 18; N = 6) or Nrxn1α KO (n = 24; N = 7) dSPNs (A) and WT (n = 20; N = 9), Nrxn1α Het (n = 15; N = 6), and Nrxn1α KO (n = 22; N = 7) iSPNs (C). (B and D) Representative confocal images of Alexa 488 fluorescence of fixed SPNs (left) and secondary dendrites from WT (top), Nrxn1α Het (middle), and Nrxn1α KO (bottom). Summary of spine density of dSPN (B) and iSPN (D) spines. Mature spines were morphologically classified as stubby and mushroom while immature include filopodia. Each point represents a neuron. Z-stacks were acquired on a confocal at 40x magnification and quantified in Image J. Data are means ± SEM; *significant difference between groups (ANOVA). See also Figure S1.

To determine whether changes in the number of striatal excitatory synapses could explain the increase in mEPSC frequency, SPNs were filled during whole-cell recordings with Alexa 488 (10uM) dye and subsequently fixed for visualization and confocal imaging (Fig.1B,D). We observed no changes in the number of spines per micrometer of dendrite onto either dSPN (WT: 13/3; 0.91 ± 0.04, Het: 16/4; 0.93 ± 0.04, KO: 16/4; 0.91 ± 0.05, ANOVA) or iSPN (WT: 14/3; 0.91 ± 0.03, Het: 12/4; 1.00 ± 0.03, KO: 7/3; 0.88 ± 0.05 spines, ANOVA) (Fig.1B,D). Further classification of spines based on morphology (stubby/mushroom/filopodia) did not reveal significant genotypic differences for either SPN subtype in both Nrxn1α heterozygote and knock-out mice (Fig.1B,D).

### Optogenetic-mediated Afferent Recruitment to Examine Synaptic Strength

The complex integrative architecture of the striatum, wherein excitatory inputs from multiple forebrain and thalamic regions make widespread, intermingled synapses with SPNs (Hunnicutt et al., 2016; Mandelbaum et al., 2019), limits the interpretability of local electrical stimulation techniques for afferent fiber recruitment. In light of this as well as evidence for distinct pools for spontaneous and evoked synaptic release (Sara et al., 2005), we employed optogenetic-mediated recruitment of specific striatal circuits to explore action-potential evoked excitatory transmission in Nrxn1α mutant mice. Prior cell type-specific monosynaptic tracing from spiny neuron and local interneuron subtypes within DMS revealed extensive connectivity from dorso-prefrontal cortical (dPFC) and parafascicular thalamic regions (Choi et al., 2019).

To isolate dPFC-striatal circuits, we transduced dPFC with AAV-DJ-hSyn-ChIEF-2a-Venus and optically evoked synaptic release in the presence of GABAR blockade. We simultaneously performed striatal field/whole-cell recordings (maximum distance 50µm) (Choi et al., 2019; Xiong et al., 2015) using the fiber volley of the field to normalize for opsin-expressing fibers (Fig.2A). For each recorded neuron, the whole-cell EPSC amplitude was plotted against the field fiber volley across increasing light intensity, with the resulting regression coefficient used as a proxy of synaptic strength (Fig.2B, see Methods for details). This approach was sensitive over a range of release probabilities, as demonstrated by the dynamic changes in regression coefficient with alterations in release probability (Fig.2C,D).

**Figure 2.**
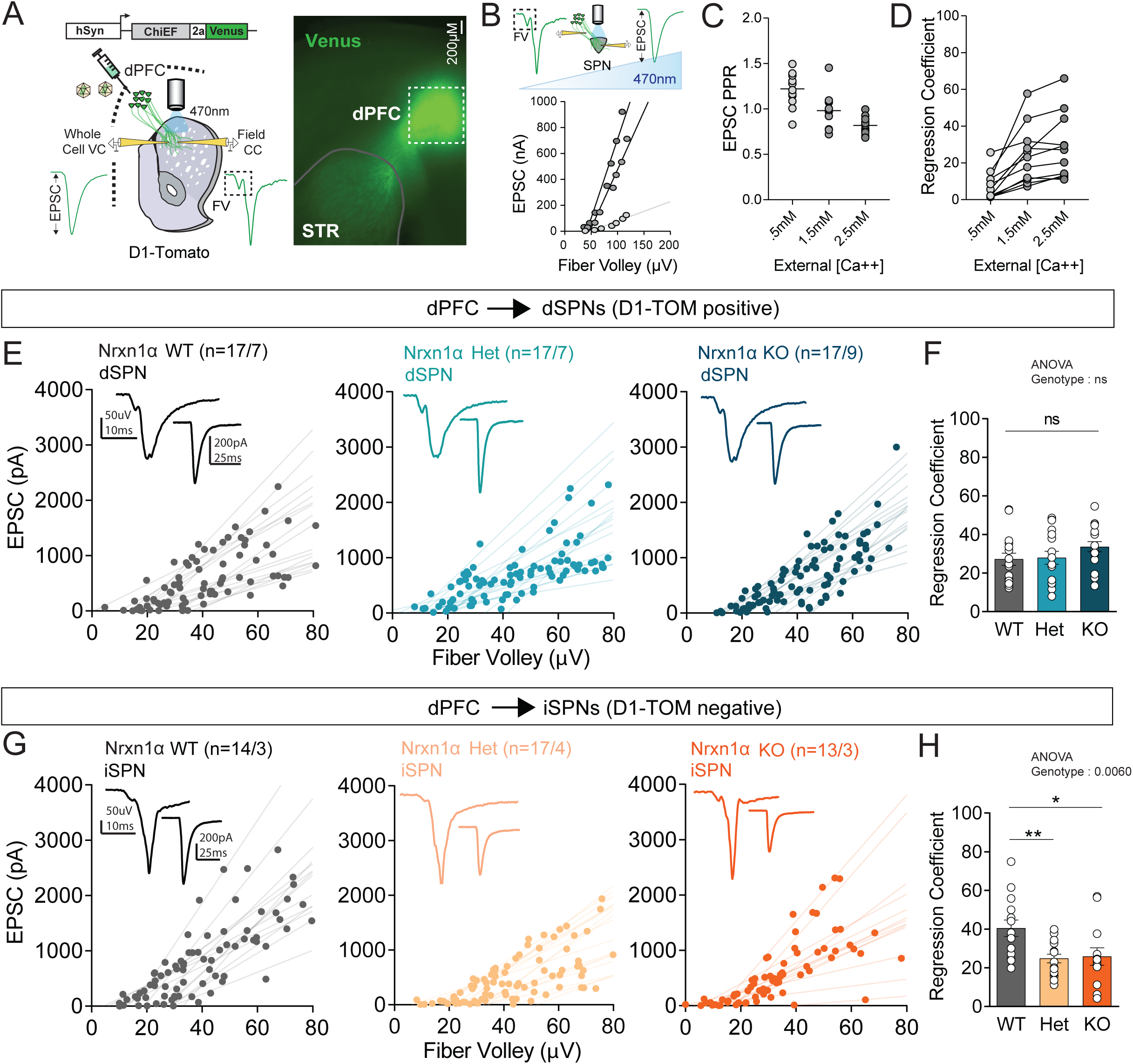
dPFC-DMS synapses exhibit a selective reduction in synaptic strength onto iSPNs in synapses in Nrxn1α Het and Nrxn1α KO animals. (A) Diagram of AAV constructs and combined whole-cell and field recording approach (left). Representative image showing ChiEF-2a-Venus expression in the dPFC (injection site, dotted white) and DMS (gray solid outline). Scale bar is 200μM. (B) Illustration of stepwise increase of 470nm LED intensity to measure a range of fiber volleys and postsynaptic EPSCs recorded in a combined whole-cell and field recording configuration (top). Representative cell depicting fitted lines from regression analyses of input output measurements across varied external calcium levels. (C) Paired-pulse ratio (50ms ISI) is sensitive to external calcium levels (.5mM, 1.5mM, 2.5mM). (D) Summary of regression coefficients from linear regression analyses performed on input output measurements. Connected lines represent a neuron across varied external calcium levels. (E and G) Plot of changes in fiber volley amplitude and EPSC across led intensities in WT (left), Nrxn1α Het (middle) and Nrxn1α KO (right) of dSPNs (E) and iSPNs (G). Inset shows representative field recording with adjacent whole-cell recording below. (F and H) Summary of regression coefficients of the linear regression analyses performed on the input output measurements of dSPNs (F) and iSPNs (H). Each point represents a neuron. Summary data are mean ± SEM, *p < 0.05. See also Figure S2.

### Target-specific Reductions in Excitatory Synaptic Strength of dPFC-striatal Circuits

We noted no changes in EPSC versus fiber volley slope for dPFC-dSPN connections in either Nrxn1α heterozygote or knock-out mice as compared to wildtype littermate controls (WT: 17/7, Het: 17/7, KO: 17/9, ANOVA) (Fig.2E, F). In contrast, we noted a significant, ∼50% reduction in the regression coefficient of both Nrxn1α heterozygote and knock-out mice onto iSPNs suggesting a decrease in synaptic strength at this connection (WT: 14/3, Het: 17/4 [p<0.01], KO: 13/3 [p<0.05], Dunnett’s test) (Fig2G,H). Changes in AP-evoked synaptic strength can arise from mutation-associated alterations in pre- or postsynaptic function. Given neurexins presynaptic localization, we probed presynaptic release via paired-pulse ratio (PPR). Optical stimulation of dPFC fibers targeting striatum revealed no changes in the PPR onto dSPNs (WT: 24/4, Het: 11/3, KO: 27/4, 2-way ANOVA), across genotypes (Fig.3A). Consistent with these data, there were no changes in the five-pulse variable frequency trains onto dSPNs across the three frequencies measured, including 10Hz, 20Hz, 50Hz (WT: 18/4, Het: 11/3, KO: 24/4, 2-way ANOVA) (Fig.3B). To make sure our inability to detect an expected increase in release probability for dPFC-dSPN connections was not a floor effect from optogenetic-associated decreases in PPR, we minimized direct terminal illumination by localizing a 50μm 470nm light spot at the corticostriatal border. PPR was not significantly higher in spot compared with full field illumination (Supp.Fig.2A-C). Nevertheless, we tested dPFC-dSPN short term dynamics in wildtype and Nrxn1α KO mice with spot illumination and reduced extracellular Ca^2+^, again detecting no genotypic differences (Supp.Fig.2D,E).

In contrast to the dPFC-dSPN pathway, we noted an increase in the PPR onto iSPNs (WT: 19/4, Het: 14/3, KO: 20/5, 2-way ANOVA) in both Nrxn1α heterozygote and knock-out mice (Fig3D). Furthermore, there was an increase in the ratios of the nth pulse/1^st^ pulse of the five-pulse variable frequency trains onto iSPNs (WT: 18/5, Het: 14/3, KO: 18/5, 2-way ANOVA (Fig.3E). To probe postsynaptic function, we measured optical NMDA/AMPA ratios observing no changes onto either cell-type (dSPNs WT: 21/4, Het: 13/3, KO: 26/4, iSPNs WT: 22/5, Het: 14/3, KO: 26/6, ANOVA) (Fig.3C,F). These results suggest a postsynaptic target-selective decrease in excitatory synaptic strength at dPFC-iSPN synapses, driven by reductions in presynaptic transmitter release probability.

**Figure 3.**
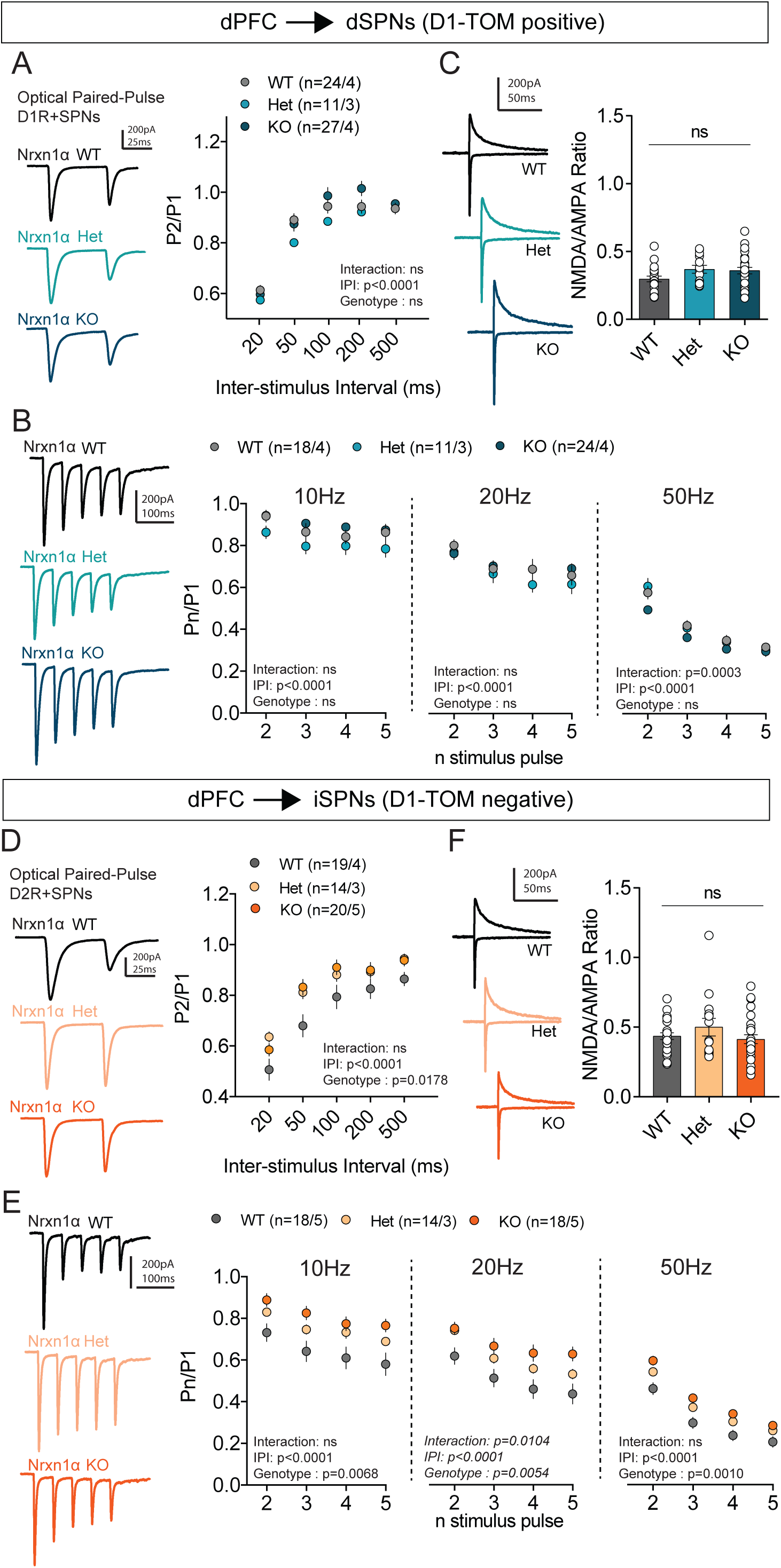
Reduced probability of release onto iSPNs at dPFC-DMS synapses in Nrxn1α Het and Nrxn1α KO animals. (A and D) Representative traces of paired-pulse response in each genotype (left; 50ms ISI) and plot of paired-pulse ratio across multiple inter-stimulus intervals (ISI) (right) of dSPNs (A) and iSPNs (D). (B and E) Representative traces of 5-pulse frequency trains (20Hz; left) and plot of frequency trains across multiple frequencies (right; 10Hz; 20Hz; 50Hz from left to right) of dSPNs (B) and iSPNs (E). (C and F) Representative traces of recordings used to measure NMDA/AMPA ratio (NMDA current is measured 50ms from stimulation). Plot of NMDA/AMPA ratio by genotype (right) of dSPNs (C) and iSPNs (F).

### Neurexin1α Deletion alters Postsynaptic Function at Thalamostriatal Synapses through a Reduction in NMDAR Current

As Nrxn1α is expressed at roughly similar levels across striatal afferent sites (Fuccillo et al., 2015), we wanted to determine whether the above reported synaptic changes were a common property across striatal projecting regions or were specific to this cortical input. We therefore targeted another region that densely innervates DMS, examining synaptic properties from the parafascicular nucleus of thalamus (Choi et al., 2019; Ellender et al., 2013; Mandelbaum et al., 2019) (Fig.4A). Using identical simultaneous field/whole-cell recordings, we measured the input-output relationship onto both SPN subtypes.

**Figure 4.**
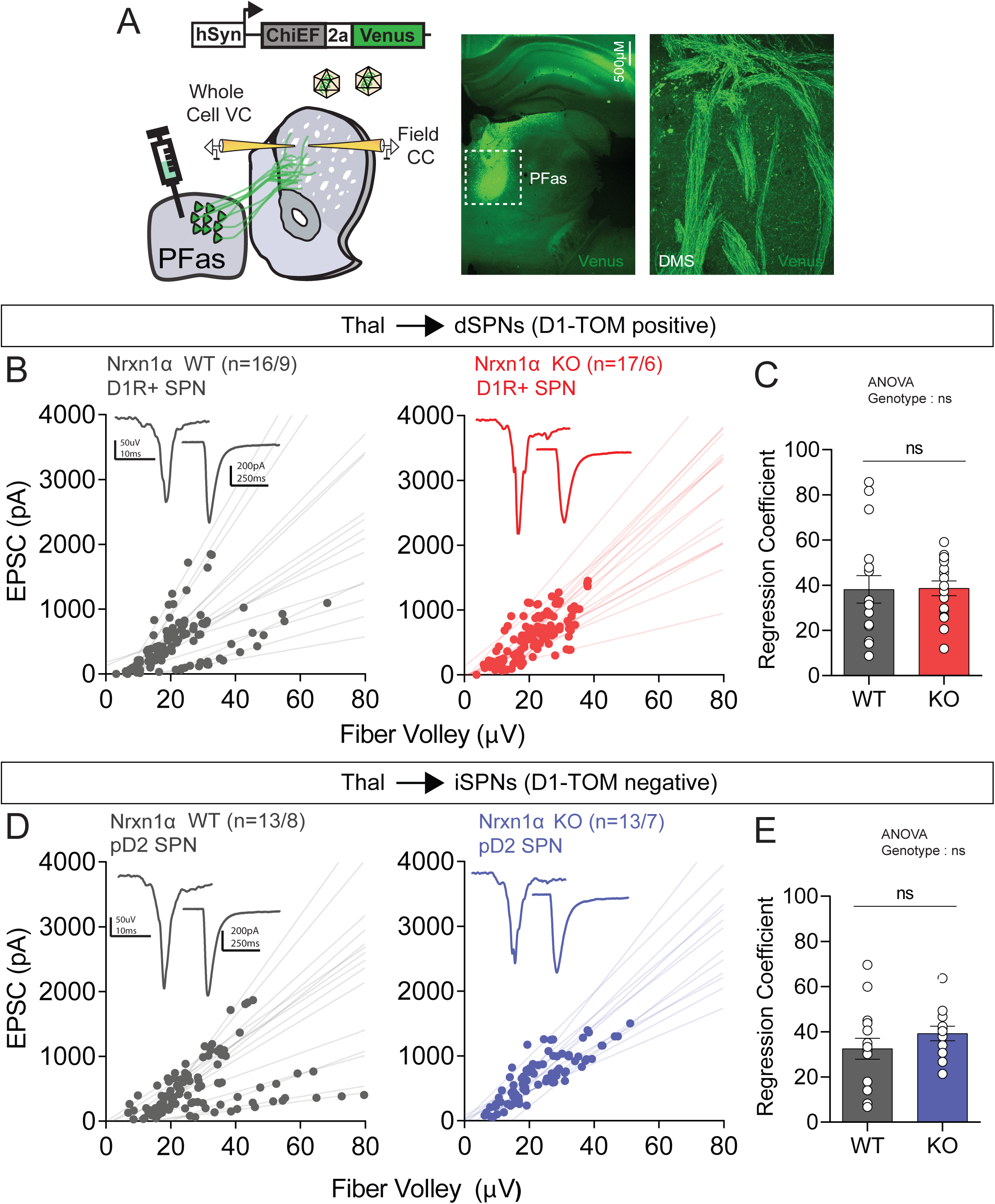
Synaptic strength is unaltered at PFas-DMS synapses in Nrxn1α KOs. (A) Experimental scheme of injection and recording sites for PFas-DMS projection (left). Representative image of ChiEF-2a-Venus expression in PFas injection site. (B and D) Plot of changes in fiber volley amplitude and EPSC across led intensities in WT (left), and Nrxn1α KO (right) of dSPNs (B) and iSPNs (D). Inset shows representative field recording with adjacent whole-cell recording below. (C and E) Summary of regression coefficients of the linear regression analyses performed on the input output measurements of dSPNs (C) and iSPNs (E). Each point represents a neuron.

Surprisingly, linear regression analysis revealed no genotypic differences for PFas excitatory synaptic connections onto either SPN subtype (dSPN, WT: 16/9, KO: 17/6, iSPN, WT: 13/8, KO: 13/7, Unpaired two-tailed t-test) (Fig.4B-E). Consistent with this result, no changes were observed in paired-pulse ratio (Fig.5A,D) or five-pulse variable frequency trains (Fig.5B,E) onto either SPN subtype. To probe for postsynaptic alterations we measured optical NMDA/AMPA ratios, detecting a significant decrease in the Nrxn1α KO mice onto both SPN subtypes (Fig.5C,F). Furthermore, we observe a significant decrease in NMDA decay at synapses onto iSPNs, suggesting alterations in subunit composition (Supp. Fig.3A-B). These results suggest that, in contrast to a presynaptic locus of action at dPFC synapses in striatum, removal of Nrxn1α alters postsynaptic function at PFas-DMS synapses onto both SPN subtypes, without impacting release probability.

**Figure 5.**
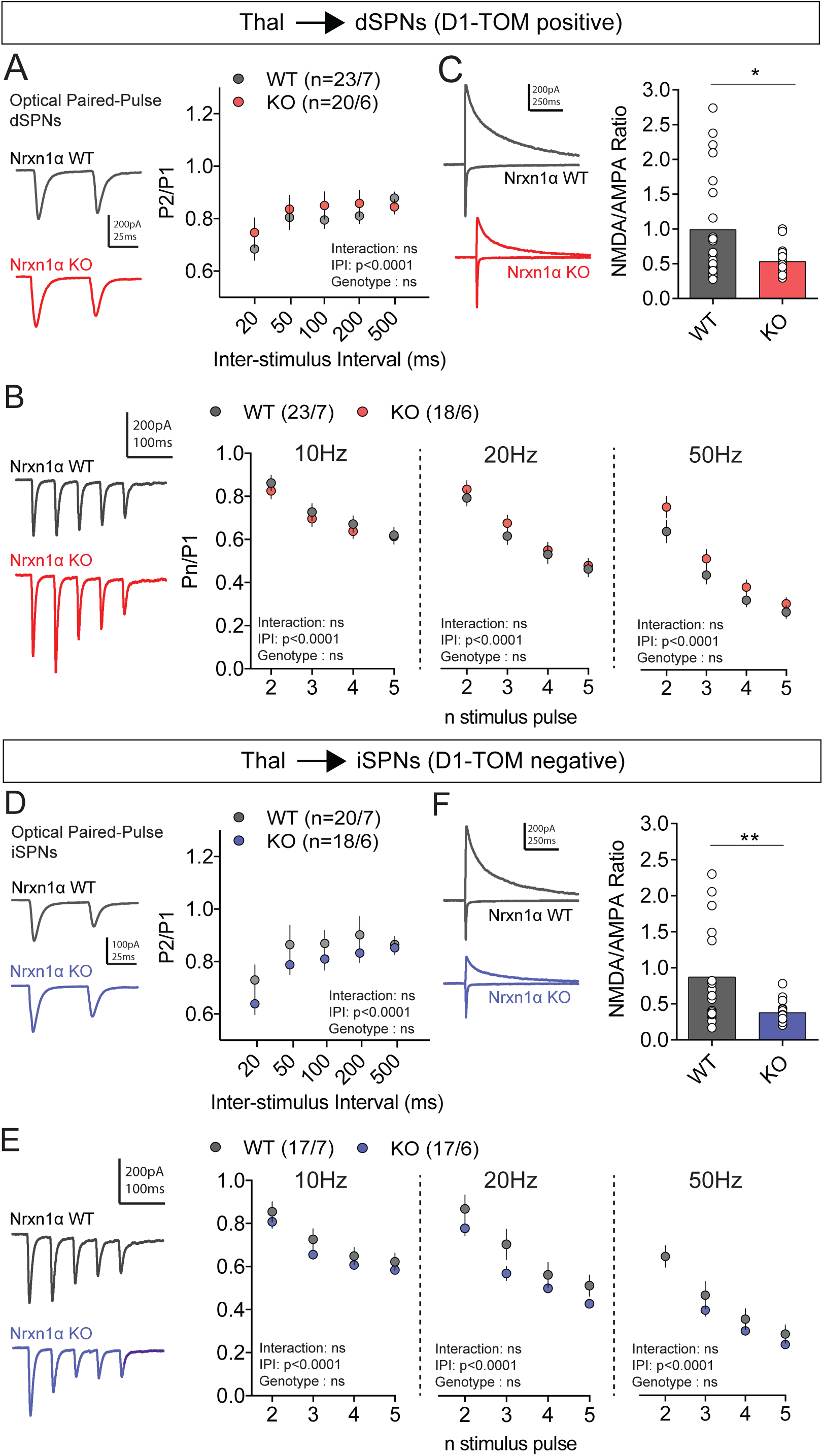
Altered NMDA/AMPA ratios at PFas-DMS projection onto both SPN subtypes in Nrxn1α KO animals. (A and D) Representative traces of paired-pulse response in each genotype (left; 50ms ISI) and plot of paired-pulse ratio across multiple ISIs (right) of dSPNs (A) and iSPNs (D). (B and E) Representative traces of 5-pulse frequency trains (20Hz; left) and plot of frequency trains across multiple frequencies (right; 10Hz; 20Hz; 50Hz from left to right) of dSPNs (B) and iSPNs (E). (C and F) Representative traces of recordings used to measure NMDA/AMPA ratio (NMDA current is measured 50ms from stimulation). Plot of NMDA/AMPA ratio by genotype (right) of dSPNs (C) and iSPNs (F). Summary data are mean ± SEM, *p < 0.05, **p < 0.01. See also Figure S3.

### Reduced Responsiveness to *In Vivo*-Modeled Input Frequencies at iSPN Corticostriatal Terminals

It remains unclear how Nrxn mutation-associated target-specific changes in presynaptic release would influence corticostriatal information transfer in the context of more intact circuits. To examine this issue, we generated *in vivo*-modeled optical stimulus patterns by using *in vivo* single-unit recordings as a mask to filter Poisson distributed cortical spike trains of frequencies typical of dPFC regions (5Hz,15Hz, 25Hz, details in Methods) (Fig.6A, Supp.Fig.4A). Experiments were performed in current-clamp, holding the neuron at -55mV to mimic the *in vivo* “up-state” membrane potential (Stern et al., 1998; Stern et al., 1997), and GABAergic inhibition was left intact. Total spiking efficiency (APs/number of optical stimuli) in the Nrxn1α KO, relative to WT, was unchanged onto dSPNs (Fig.6B) and decreased onto iSPNs (Fig.6D). Input-output coupling deficiencies were observed at dPFC-iSPN connections across all tested input frequency domains (Fig.6C,E). These results were not the result of biases in initial EPSP amplitude and did not depend on recording duration (Supp.Fig.4B-E). Together these findings suggest that the target cell-specific reductions in release probability observed for dPFC-iSPN circuits have significant impact on the fidelity of corticostriatal connectivity.

**Figure 6.**
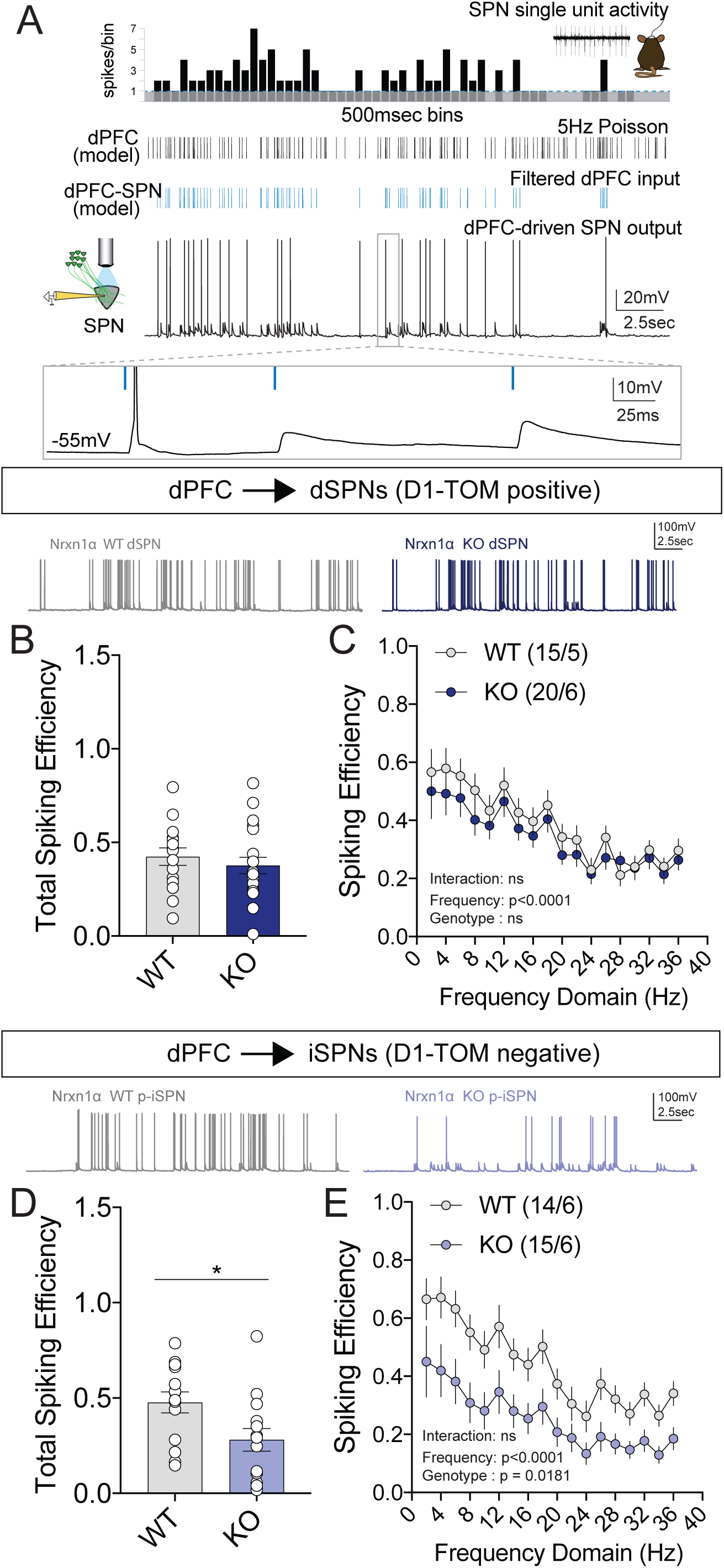
Reduced synaptic efficacy across a broad range of naturalistic input frequencies. (A) Local inhibitory circuit remained intact for current-clamp recordings to permit a global view of synaptic efficacy. Optical stimulus patterns were modeled after SPN firing in *in vivo* single-unit recordings. (Top) Thresholded (>2Hz, above dotted line) *in vivo* SPN activity is used as a mask for Poisson distributed cortical spike trains (black) to generate optogenetic pattern (blue). (Bottom) SPN spiking output to “modeled” dPFC-SPN inputs. (B,D) Overall spiking efficiency for each dSPN and iSPN (D) recorded measured as the number of action potentials for a given optical pattern. (C,E) Spiking efficiency across local frequencies represented in the filtered dPFC input stimulus for current-clamp recordings done in dSPNs and (E) iSPNs. Spiking efficiency = action potentials/number of optical inputs. Summary data are mean ± SEM, *p < 0.05. See also Figure S4.

**Figure 7.**
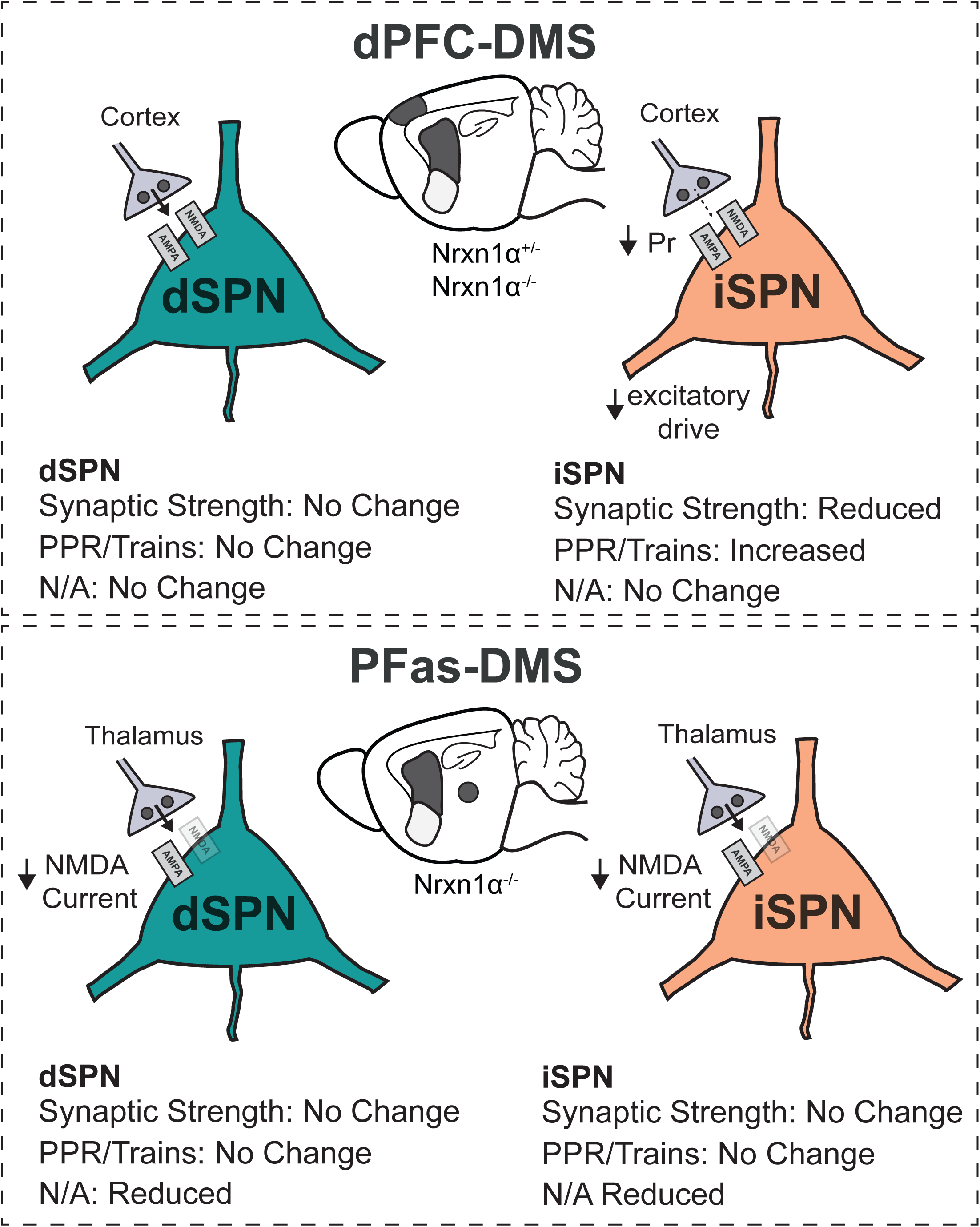
Divergent synaptic phenotypes at dPFC-DMS and PFas-DMS synapses in Nrxn1α mutants. (Top) Optical evoked synaptic measurements at dPFC-DMS synapses reveal no changes onto dSPNs (left), however there was a decrease in synaptic strength onto iSPNs with an increase in PPR/Trains in both Nrxn1α heterozygote and knockout animals. (Bottom) Probing PFas-DMS synapses using optical measurements we detect changes no changes in synaptic strength or probability of release measurements, however, we find reductions in NMDA/AMPA ratio onto both SPN subtypes in Nrxn1α KO animals.

## Discussion

Neurexins are presynaptic cell-adhesion molecules exhibiting diverse synaptic functions dependent on cell-type and neural circuit (Etherton et al., 2009; Sudhof, 2008, 2018). Here, we show that Nrxn1α differentially regulates glutamatergic excitatory synaptic transmission at dPFC and PFas inputs to DMS. In both Nrxn1α heterozygous and homozygous animals we observed reductions in probability of release onto iSPNs, but not dSPNs, at dorsal prefrontal cortical inputs to DMS. We demonstrated that reduced synaptic efficacy onto iSPNs is not a common feature of all striatal targeting inputs, as PFas inputs have normal release probability, while exhibiting altered postsynaptic NMDAR function. These findings provide further evidence that synaptic transmission regulated by Nrxn1α is input and target specific, illustrating for the first time its role within striatal circuits.

### Challenges of Studying Presynaptic Neurexin1α Function in Striatal Circuits

Advances in understanding the role of cell-adhesion molecules located pre-synaptically have been hampered by technical challenges. Earlier studies employing electrical stimulation were limited by the heterogeneity of stimulated inputs as well as the substantial recruitment of systems that modulate neuronal function (Cavaccini et al., 2018; Kreitzer and Malenka, 2007; Shen et al., 2008). To gain access to specific striatal circuits, we utilized optogenetics, permitting light-mediated selective recruitment of either dPFC or PFas striatal inputs through the expression of a channelrhodopsin variant, ChiEF-2a-Venus (Lin et al., 2009). To normalize for differences in viral expression we employed a combined field and whole cell recording approach (Choi et al., 2019). The optically-evoked field recorded in striatum is composed of two negative components (N1, N2) (Kupferschmidt et al., 2017), with the first negativity being largely composed of presynaptic opsin-expressing fibers. We used this signal as a normalization measurement for afferent recruitment when measuring currents onto neighboring neurons. This approach provides an important step forward for understanding long-range synaptic connectivity, circumventing limitations regarding efficacy of viral transduction (Chuhma et al., 2011; Lenz and Lobo, 2013; Miesenbock, 2011). Furthermore, it permits more targeted access to specific neuronal populations than that provided by broadly-expressed Cre lines (Rbp4-Cre, Thy1-Cre) previously employed to measure synaptic strength. A limitation of this technique is the requirement for relatively dense targeting by the afferent population, which is necessary to observe a clear, reliable optical fiber volley.

### Convergent Physiological Phenotypes in Nrxn1α -/+ and Nrxn1α -/- Mice

Genes encoding proteins that support the formation and maintenance of synapses are significantly associated with ASD and schizophrenia (Chang et al., 2015). Disease-associated mutations in *Nrxn1* are commonly found as heterozygous copy number variations (CNVs) (Autism Genome Project et al., 2007; Huang et al., 2017; Marshall et al., 2017). Behavioral studies using mice with either heterozygous or homozygous Nrxn1α deletions reveal non-overlapping phenotypes, with KOs exhibiting elevated anxiety levels, increased aggressive behaviors, nest building impairments, and altered social behaviors not observed in heterozygotes (Grayton et al., 2013). Despite this phenotypic divergence, Nrxn1α homozygous deletions have been the central focus of physiological analyses in rodents (Etherton et al., 2009). Studies in human embryonic stem cells highlight the relevance of heterozygous deletions, revealing substantial impairments in excitatory synaptic transmission (Pak et al., 2015). It nevertheless remains unclear whether haploinsufficiency of Nrxn1α disrupts neuronal physiology in disease relevant neural circuits. Our data clearly demonstrates synaptic impairments within dPFC-iSPN circuits of roughly similar magnitude in both Nrxn1α heterozygous and homozygous mice. These data contrast with the absence of excitatory synaptic phenotypes in cortical primary neuronal cultures from Nrxn1α heterozygous and homozygous mice, highlighting the diversity of Neurexin function dependent on synaptic context (Anderson et al., 2015; Aoto et al., 2013).

### Nrxn1α Regulates Synaptic Transmission at dPFC-DMS Synapses in a Cell-type Specific Manner

An interesting aspect of our study is the specificity of altered dPFC glutamatergic transmission dependent on target SPN subtype. Mice with either heterozygous or homozygous deletion of Nrxn1α were observed to have decreased synaptic strength onto striatal iSPNs. While generally consistent with reductions in input-output strength observed at CA1 hippocampal synapses (Etherton et al., 2009), here we note the reduction in synaptic strength onto striatal iSPNs likely results from decreased release probability, as demonstrated by elevated paired pulse ratios and reductions in synaptic depression upon high-frequency stimulation. One surprising aspect of this phenotype is that cortico-striatal afferents do not show significant target-neuron specificity, as evidenced both by cell type-specific retrograde rabies labeling (Wall et al., 2013) and single axonal tracing (Doig et al., 2010). While this lack of anatomical specificity contrasts with our observed synaptic phenotypes, it is possible that molecular diversity of Neurexin signaling exists at the individual spine level, an interesting hypothesis awaiting higher resolution transcriptomic and proteomic approaches.

In addition to this puzzling target specificity, the exact pathways by which Nrxn1α disruption leads to physiological dysfunction remain unclear. Potential mechanisms include changes in the organization of synaptic release machinery or alterations in GPCR-coupled signaling pathways that modulate neurotransmitter release. Previous work showed that presynaptic β-Neurexins inhibit tonic endocannabinoid synthesis, thereby maintaining hippocampal CA1-subicular synaptic strength (Anderson et al., 2015). Furthermore, postsynaptic neuroligin-3 regulates tonic endocannabinoid signaling to constrain local basket cell inhibition (Foldy et al., 2013). Given these data, it is tempting to speculate that our observed decrease in release probability and synaptic strength in dPFC-iSPN circuits of Nrxn1α mutants emerges from dysregulation of eCB-mediated control. This hypothesis may also explain the observed input (dPFC but not PFas) and target (iSPN but not dSPN) specificity insofar as – (1) higher CB1R expression patterns in cortex than thalamus restrict eCB-mediated depression to corticostriatal synapses (Wu et al., 2015) and (2) prior implication of specific 2-AG pathway dysfunction in β-Neurexin mutants (Anderson et al., 2015) may bias depression to the iSPN subtype (Ade and Lovinger, 2007; Giuffrida et al., 1999).

### Nrxn1α Maintains NMDAR Function at PFas-DMS Synapses onto both SPN Subtypes

Given the broad expression of *Nrxn1* throughout striatal-targeting projections, we asked whether alterations in synaptic efficacy is unique to dPFC-striatal circuits. We employed similar optogenetic methods to probe parafascicular thalamic inputs to the dorsal striatum but did not detect changes in synaptic strength or release probability. Interestingly, in contrast to dPFC-DMS synapses, NMDA/AMPA ratios at PFas-DMS synapses were reduced onto both SPN subtypes in the KO mice. We hypothesize that this reduced ratio results from a decrease in NMDAR currents for several reasons – (1) increases in AMPAR current should have increased the input/output slope measurements, (2) there was no evidence for changes in the amplitude of excitatory mEPSCs (although PFas neurons only represent a fraction of the total inputs recorded with this technique) and (3) triple KO of α-neurexins were observed to reduce NMDAR function (Kattenstroth et al., 2004). While reduced synaptic NMDAR content may reflect a decrease in postsynaptic recruitment of Neuroligin1, which is known to mediate clustering of NMDARs (Blundell et al., 2010; Budreck et al., 2013; Wu et al., 2019), the lack of this phenotype at dPFC-striatal synapses remains unclear. It is worth considering the disconnect between our original spontaneous transmission data and the complexity of the dPFC and PFas synaptic phenotypes observed with pathway-specific optical recruitment. Whether these discrepancies arise from the diverse nature of excitatory afferent projections into DMS or represent the parallel actions of spontaneous and action-potential-evoked synaptic transmission pathways is yet unclear (Kavalali, 2015).

### Changes to Striatal Circuit Output and its Implications

Cortical and thalamic regions exhibit topographic projections to discrete striatal compartments, where their control of SPN activity and local inhibitory circuits is an initial step in the generation of motor output. We extended our initial discovery of reduced synaptic strength in dPFC-iSPNs circuits of Nrxn1α -/- mice by probing the fidelity of spike production for dPFC inputs modeled on *in vivo* activity patterns. Consistent with our target cell-specific effects, we found that dPFC-iSPN but not dPFC-dSPN circuits exhibited reduced total spiking efficiency across a range of frequency domains. This selective decrease in cortical-iSPN excitatory drive is expected to promote disinhibition of thalamic structures projecting back to cortex. It is an interesting future question as to whether this thalamic disinhibition can explain both simple motor phenotypes (e.g. hyperactivity, aggression) (Etherton et al., 2009; Grayton et al., 2013) and more complex reward processing deficits observed in mice with Nrxn1α mutations (Alabi et al., 2019). It is worth noting a recent study of the *Tsc1* deletion ASD mouse model, which exhibited increased corticostriatal connectivity selectively onto dSPNs, with synaptic strength unperturbed at thalamostriatal synapses (Benthall et al., 2018). Together with our data, this raises the possibility that disinhibition of BG-targeted thalamus is a common circuit disruption for ASD pathophysiology with clear implications for downstream cortical activity.

Here we demonstrated that the loss of Nrxn1α has differential effects within striatal circuits, extending the literature suggesting imbalances in basal ganglia activity as a common perturbation in models of neurodevelopmental disorders. Our data highlight the challenges that context-dependent gene function creates for understanding molecular contributions to brain and behavioral pathology. Diversity of inputs is even apparent for cortical sub-regions, which display profound heterogeneity in anatomical and functional connectivity to striatum as well as divergent synaptic phenotypes in response to loss of the postsynaptic scaffolding molecule SAPAP3, a gene linked to neuropsychiatric disorders (Corbit et al., 2019). Given that genes related to synaptic function are implicated in neuropsychiatric disease, detailed circuit-specific studies *in vitro* and *in vivo* are needed in elucidating their context-specific functions, thereby gaining a better understanding of how synaptic dysregulation can contribute to pathogenesis.

### Experimental Procedures

All experiments were conducted in accordance with the National Institutes of Health Guidelines for the Use of Animals, and all procedures approved by the Institutional Animal Care and Use Committee of the University of Pennsylvania (Protocol: 805643). Animals were kept on a 12:12 light-dark cycle and provided food and water *ad libitum*.

### Animals

Constitutive Nrxn1α KO mice were obtained from the Südhof lab (Stanford University) (Geppert et al., 1998). To yield mice for breeding, founders were crossed onto C57BL/6 (Jackson Laboratory) generating Nrxn1α ^+/-^ animals. To identify direct pathway and putative indirect pathway neurons, Nrxn1α ^+/-^ animals were subsequently crossed onto the Drd1a-tdTomato BAC transgenic line. Breeders for experimental animals were male and female Nrxn1α ^+/-^ animals with one breeder also hemizygous for D1Tom^+^. Offspring were weaned at P21 and separated by sex in cages of 2-5 animals of mixed genotypes.

### Stereotaxic Surgeries

Adult male mice were between the ages of 3-5 weeks were injected into regions of interest. Viral injections were performed on a stereotaxic frame (Kopf Instruments, Model 1900) under isoflurane anesthesia (3% for induction; 1-2% during surgery). Sterile surgical technique was used, removing fur over the skull with a depilatory cream, and applying 70% isopropyl alcohol and betadine. Prior to creating an incision in the scalp to access the skull, a local anesthetic, lidocaine, was given subcutaneously. Small (0.5mm) holes were drilled and a pulled glass capillary needle was carefully lowered into the brain. Coordinates relative to bregma used for the regions of interest include dorsal prefrontal cortex (AP: +1.9mm, ML: +/- 0.3mm, DV: -1.4mm) and parafasicular nucleus of thalamus (AP: -2.2, ML: +/- 0.75, DV: -3.7). Bilateral injections of AAV-DJ-hSyn-ChiEF-2a-Venus (volume: 300nl) were infused 100nl/min using a microinfusion pump (Harvard Apparatus, #70-3007). The capillary glass was kept in the target region for 5 minutes after viral infusion was complete to prevent backflow, brought up .2mm, and after an additional 5 minutes was slowly brought up out of the brain. Mice were sutured with non-absorbable monomid nylon sutures (Stoelting), subcutaneously injected with carprofen for inflammation, and given 3 weeks (for dPFC) or 4-5 weeks (for PFas) to allow for recovery and viral expression. Prior to acute slice recording, target site injections are confirmed by examining viral expression in the anterior slices (for dPFC) and posterior slices (for PFas).

### Electrophysiology

Mice were deeply anesthetized and perfused transcardially with ice-cold ACSF (pH 7.3-7.4) containing (in mM): 124 NaCl, 2.5 KCl, 1.2 NaH_2_PO_4_, 24 NaHCO_3_, 5 HEPES, 12.5 glucose, 1.3 MgSO_4_, 7H_2_O, 2.5 CaCl_2_. The brain was rapidly removed, and coronal sections (250uM) unless otherwise indicated, were cut on a vibratome (VT1200s, Leica). Slices were incubated in a holding chamber for 12-15 minutes at 32-34°C in a NMDG-based recovery solution (pH 7.3-7.4, pH adjusted with HCl) (in mM): 92 NMDG, 2.5 KCl, 1.2 NaH_2_PO_4_, 30 NaHCO_3_, 20 HEPES, 25 glucose, 5 sodium ascorbate, 2 thiourea, 3 sodium pyruvate, 10 MgSO_4_, 7H_2_O, 0.5 CaCl_2_. Osmolarity for the NMDG-based solution and ACSF was kept between 300-310 mOsm. Following incubation, slices were moved to a second holding chamber containing ACSF at room temperature (20-22°C) for at least 1 hr. prior to recording. For recording, slices are transferred to the recording chamber (Scientifica) fully submerged in oxygenated (95% O_2_, 5% CO_2_) ACSF at a perfusion rate of 1.4-1.6 mL/min, bath temperature of 29-30°C, and secured using a slice anchor (Warner Instruments). Electrophysiology data were acquired using custom-built Recording Artist software version (Rick Gerkin), Igor Pro 6.37 (Wavemetrics). All recordings were sampled at 20kHz, filtered at 2.8kHz except for the *in vivo* modeled trains that were sampled at 10kHz.

### Intracellular Recording

Striatal SPNs were visualized using differential interference contrast (DIC) video microscopy on an upright microscope (Olympus, BX51). Somatic whole-cell recordings were performed using borosilicate glass (World Precision Instruments, TW150-3) that had a tip resistance of 3-5 MΩ, filled with cesium-based internal for voltage-clamp recordings (in mM): 115 CsMeSO_3_, 20 CsCl, 10 HEPES, 0.6 EGTA, 2.5 MgCl, 10 Na-Phosphocreatine, 4 Na-ATP, 4 Na-GTP, 0.1 Spermine, 1 QX-314 (pH adjusted to 7.3-7.4 with CsOH) or potassium-based internal (in mM): 140 K-gluconate, 5 KCl, 0.2 EGTA, 2 MgCl_2_, 10 HEPES, 4 Mg-ATP, 0.3 Na-GTP, 10 Na-Phosphocreatine (pH adjusted to 7.3-7.4 with KOH). For the striatal field recordings, a pipette was filled with ACSF and had an access resistance between 0.8-1.2 MΩ. Voltage-clamp recordings were done holding the cell at a membrane potential of -70mV (unless otherwise indicated) using a MultiClamp 700B patch-clamp amplifier (Molecular Devices). Input resistance and access resistance were noted subsequently following membrane break-in, and cells with R_A_>25 MΩ or R_I_>300 MΩ were discarded from further analysis.

For optogenetics, we used full-field 470nm illumination through a 40x objective (Olympus, 0.8NA water immersion) with a pulse width of 1ms. Blue light from a collimated LED illuminator (CoolLED, PE-300). To rule out somatic opsin contamination, an elongated light pulse was given following the last cell recorded in the slice. Optical-evoked voltage-clamp recordings were performed in the presence of picrotoxin (100uM), a GABA_A_ antagonist. AMPAR/NMDAR ratios were determined by comparing peak amplitude of averaged AMPAR EPSCs at -70mV (15 traces), with an averaged amplitude of EPSCs recorded at +40mV (15 traces), 50ms after optical stimulation. LED intensities ranged from 0.042-0.543 mW/mm^2^ during the optical input output measurements.

All miniature synaptic currents were recorded in the presence of tetrodotoxin (1μM). For excitatory currents we used the cesium-based internal described above, however, for inhibitory currents we used an internal with high chloride to maximize IPSC amplitude (in mM): 135 CsCl, 10 HEPES, 0.6 EGTA, 2.5 MgCl, 10 Na-Phosphocreatine, 4 Na-ATP, 0.3 Na-GTP, 0.1 spermine, 1 QX-314.

### Data and Statistical Analysis

Statistics was performed with Graphpad Prism v7.0. All data are presented as the mean ± SEM, with N referring to the number of animals and n to the number of cells. See supplementary tables for statistical values. Neuronal spiking was detected in NeuroMatic v.3.0 (Rothman JS, Frontiers 2018) where further analysis was then performed in custom MATLAB (Mathworks) scripts. Miniature postsynaptic currents were measured in Minianalysis (Synaptosoft). Anatomical data examining spines were measured using script written for ImageJ (NIH) with investigator blinded to genotype. Z stacks were taken on a Leica SP5II confocal microscope (fluorescent). Image manipulation and figure generation were performed in Image J, Adobe Illustrator, and Adobe Photoshop.

## Acknowledgements

This work was supported by grants from the Howard Hughes Gilliam Fellowship and Behavioral and Cognitive Training Grant (T32-MH017168) to M.F.D, NIMH (R01MH115030) and IDDRC Grant from the Children’s Hospital of Philadelphia to M.V.F. We thank Nathan Henderson for his assistance with ImageJ, Alexandria Cowell for assistance with animal husbandry, Manivannan Subramaniyan with assistance in MATLAB coding, and Alexxai Kravitz for in vivo recording data that was used to design optical patterns. We also thank Kyuhyun Choi and members of the lab for insightful feedback on the manuscript draft and project.

## Author Contributions

Conceptualization, M.F.D. and M.V.F.; Methodology, M.F.D. and M.V.F.; Formal Analysis, M.F.D.; Investigation, M.F.D.; Writing – Original Draft, M.F.D.; Writing – Review and Editing, M.F.D. and M.V.F. Visualization, M.F.D. and M.V.F.; Supervision, M.F.D and M.V.F. Funding Acquisition, M.V.F.

## Figure Legends

**Table S1.** Student’s t-test values, related to Figures 4, 5, 6, and S3.

| Figure | Test | t and p values |
| --- | --- | --- |
| 4C | Two-tailed Student's t-test | $t_{31} = 0.07521$, $p = 0.9405$ |
| 4E | Two-tailed Student's t-test | $t_{26} = 1.157$, $p = 0.2578$ |
| 5C | Two-tailed Student's t-test | $t_{39} = 2.568$, $p = 0.0142$ |
| 5F | Two-tailed Student's t-test | $t_{34} = 3.101$, $p = 0.0039$ |
| 6B | Two-tailed Student's t-test | $t_{33} = 0.7286$, $p = 0.4714$ |
| 6D | Two-tailed Student's t-test | $t_{27} = 2.411$, $p = 0.0230$ |
| S3A | Two-tailed Student's t-test | $t_{36} = 0.6048$, $p = 0.5491$ |
| S3B | Two-tailed Student's t-test | $t_{32} = 2.632$, $p = 0.0129$ |

**Table S2.**
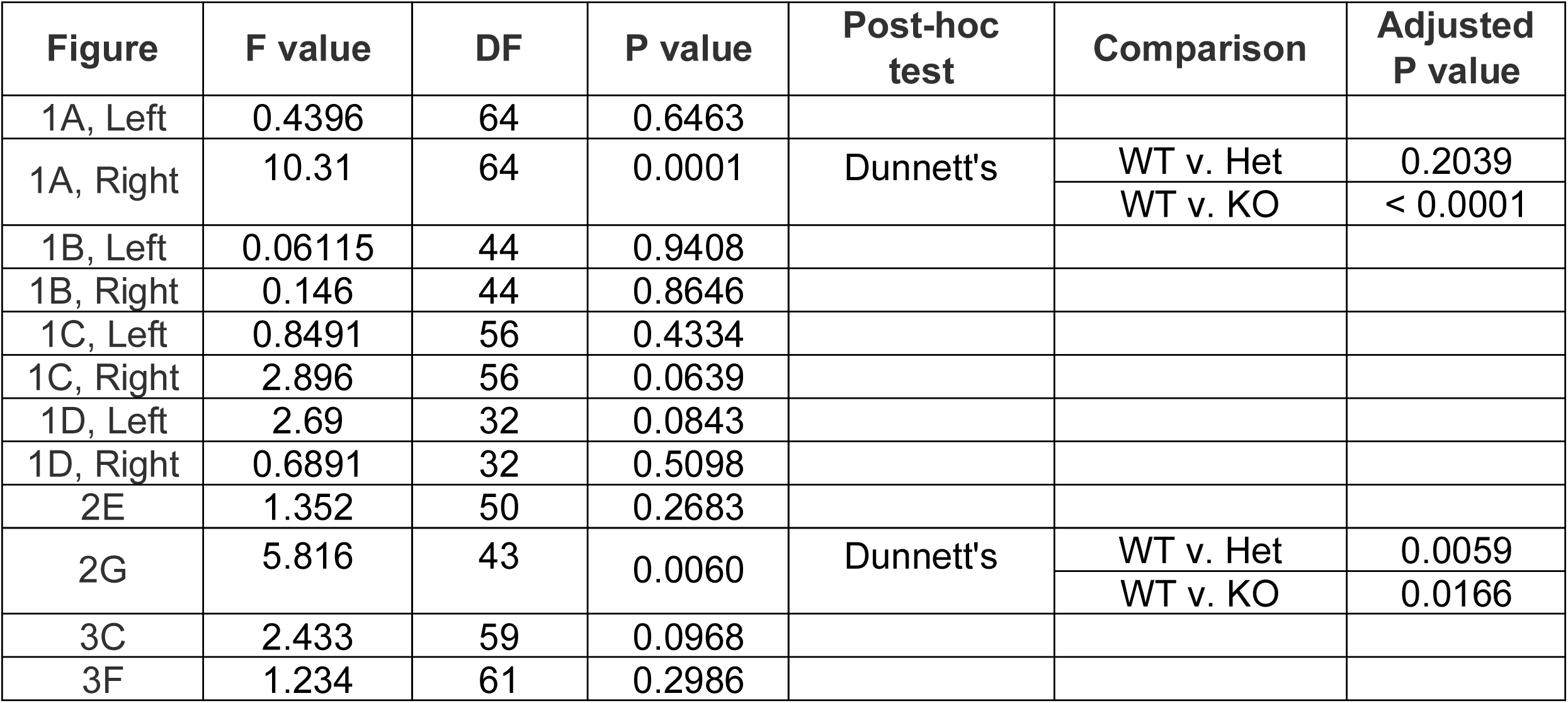
One-way ANOVA values, related to Figures 1, 2, and 3.

**Table S3.** Two-way ANOVA values, related to Figures 3, 5, and 6.

| Figure | Test | Source of Variation | F value | Comparison | DF | P value |
| --- | --- | --- | --- | --- | --- | --- |
| 3A | ANOVA | Interaction | 1.614 |  | 8 | 0.1215 |
|  |  | ISI | 144.3 |  | 4 | < 0.0001 |
|  |  | Genotype | 1.46 |  | 2 | 0.2405 |
| 3B (10Hz) | ANOVA | Interaction | 0.8324 |  | 6 | 0.5466 |
|  |  | Pulse # | 17.56 |  | 3 | < 0.0001 |
|  |  | Genotype | 1.57 |  | 2 | 0.2182 |
| 3B (20Hz) | ANOVA | Interaction | 2.026 |  | 6 | 0.0655 |
|  |  | Pulse # | 36.11 |  | 3 | < 0.0001 |
|  |  | Genotype | 0.5166 |  | 2 | 0.5997 |
| 3B (50Hz) | ANOVA | Interaction | 4.479 |  | 6 | 0.0003 |
|  |  | Pulse # | 325 |  | 3 | < 0.0001 |
|  |  | Genotype | 1.465 |  | 2 | 0.2407 |
| 3D | ANOVA | Interaction | 1.594 |  | 8 | 0.1286 |
|  |  | ISI | 175.1 |  | 4 | < 0.0001 |
|  |  | Genotype | 4.372 |  | 2 | 0.0178 |
|  | Dunnett's |  |  | ISI 20, WT v. Het | 28.13 | 0.2012 |
|  |  |  |  | ISI 20, WT v. KO | 26.08 | 0.0187 |
|  |  |  |  | ISI 50, WT v. Het | 29.84 | 0.0160 |
|  |  |  |  | ISI 50, WT v. KO | 28.88 | 0.0298 |
|  |  |  |  | ISI 100, WT v. Het | 28.83 | 0.0811 |
|  |  |  |  | ISI 100, WT v. KO | 28.39 | 0.2015 |
|  |  |  |  | ISI 200, WT v. Het | 30.49 | 0.2436 |
|  |  |  |  | ISI 200, WT v. KO | 30.7 | 0.2598 |
|  |  |  |  | ISI 500, WT v. Het | 31 | 0.0771 |
|  |  |  |  | ISI 500, WT v. KO | 32.46 | 0.0354 |
| 3E (10Hz) | ANOVA | Interaction | 0.8491 |  | 6 | 0.5342 |
|  |  | Pulse # | 87.51 |  | 3 | < 0.0001 |
|  |  | Genotype | 5.553 |  | 2 | 0.0068 |
|  | Dunnett's |  |  | Pulse 2/1, WT v. Het | 29.22 | 0.0113 |
|  |  |  |  | Pulse 2/1, WT v. KO | 23.27 | 0.0838 |
|  |  |  |  | Pulse 3/1, WT v. Het | 27.95 | 0.0086 |
|  |  |  |  | Pulse 3/1 WT v. KO | 23.99 | 0.1225 |
|  |  |  |  | Pulse 4/1, WT v. Het | 28.81 | 0.0255 |
|  |  |  |  | Pulse 4/1, WT v. KO | 26.65 | 0.0901 |
|  |  |  |  | Pulse 5/1, WT v. Het | 26.01 | 0.0124 |
|  |  |  |  | Pulse 5/1, WT v. KO | 21.33 | 0.1328 |
| 3E (20Hz) | ANOVA | Interaction | 2.605 |  | 6 | 0.0200 |
|  |  | Pulse # | 133.4 |  | 3 | < 0.0001 |
|  |  | Genotype | 5.845 |  | 2 | 0.0054 |
|  | Dunnett's |  |  | Pulse 2/1, WT v. Het | 29.36 | 0.0234 |
|  |  |  |  | Pulse 2/1, WT v. KO | 21.91 | 0.0161 |
|  |  |  |  | Pulse 3/1, WT v. Het | 29.99 | 0.0197 |
|  |  |  |  | Pulse 3/1 WT v. KO | 27.02 | 0.1101 |
|  |  |  |  | Pulse 4/1, WT v. Het | 30 | 0.0149 |
|  |  |  |  | Pulse 4/1, WT v. KO | 24.76 | 0.1149 |
|  |  |  |  | Pulse 5/1, WT v. Het | 28.97 | 0.0053 |
|  |  |  |  | Pulse 5/1, WT v. KO | 24.63 | 0.1525 |
| 3E (50Hz) | ANOVA | Interaction | 1.221 |  | 6 | 0.2991 |
|  |  | Pulse | 427 |  | 3 | < 0.0001 |
|  |  | Genotype | 8.012 |  | 2 | 0.0010 |
|  | Dunnett's |  |  | Pulse 2/1, WT v. Het | 29.49 | 0.0016 |
|  |  |  |  | Pulse 2/1, WT v. KO | 28.16 | 0.0418 |
|  |  |  |  | Pulse 3/1, WT v. Het | 29.77 | 0.0028 |
|  |  |  |  | Pulse 3/1 WT v. KO | 21.99 | 0.0297 |
|  |  |  |  | Pulse 4/1, WT v. Het | 29.7 | 0.0057 |
|  |  |  |  | Pulse 4/1, WT v. KO | 25.25 | 0.0512 |
|  |  |  |  | Pulse 5/1, WT v. Het | 29.89 | 0.0377 |
|  |  |  |  | Pulse 5/1, WT v. KO | 27.98 | 0.1281 |
| 5A | ANOVA | Interaction | 1.373 |  | 4 | 0.2456 |
|  |  | ISI | 11.36 |  | 4 | < 0.0001 |
|  |  | Genotype | 0.4059 |  | 1 | 0.5276 |
| 5B (10Hz) | ANOVA | Interaction | 0.6332 |  | 3 | 0.5950 |
|  |  | ISI | 120.9 |  | 3 | < 0.0001 |
|  |  | Genotype | 0.2746 |  | 1 | 0.6032 |
| 5B (20Hz) | ANOVA | Interaction | 0.8874 |  | 3 | 0.4499 |
|  |  | ISI | 200 |  | 3 | < 0.0001 |
|  |  | Genotype | 0.4443 |  | 1 | 0.5090 |
| 5B (50Hz) | ANOVA | Interaction | 1.962 |  | 3 | 0.1235 |
|  |  | ISI | 258.7 |  | 3 | < 0.0001 |
|  |  | Genotype | 1.891 |  | 1 | 0.1769 |
| 5D | ANOVA | Interaction | 0.543 |  | 4 | 0.7044 |
|  |  | ISI | 13.53 |  | 4 | < 0.0001 |
|  |  | Genotype | 0.9886 |  | 1 | 0.3267 |
| 5E (10Hz) | ANOVA | Interaction | 0.632 |  | 3 | 0.5961 |
|  |  | Pulse # | 119.3 |  | 3 | < 0.0001 |
|  |  | Genotype | 1.098 |  | 1 | 0.3026 |
| 5E (20Hz) | ANOVA | Interaction | 1.953 |  | 3 | 0.1262 |
|  |  | Pulse # | 201.2 |  | 3 | < 0.0001 |
|  |  | Genotype | 2.062 |  | 1 | 0.1607 |
| 5E (50Hz) | ANOVA | Interaction | 2.032 |  | 3 | 0.1146 |
|  |  | Pulse # | 249.9 |  | 3 | < 0.0001 |
|  |  | Genotype | 0.5889 |  | 1 | 0.4485 |
| 6C | ANOVA | Interaction | 0.6695 |  | 17 | 0.8340 |
|  |  | Frequency | 27.32 |  | 17 | < 0.0001 |
|  |  | Genotype | 0.6145 |  | 1 | 0.4387 |
| 6E | ANOVA | Interaction | 1.074 |  | 17 | 0.3764 |
|  |  | Frequency | 32.09 |  | 17 | < 0.0001 |
|  |  | Genotype | 6.328 |  | 1 | 0.0181 |

## References

Ade, K.K., and Lovinger, D.M. (2007). Anandamide regulates postnatal development of long-term synaptic plasticity in the rat dorsolateral striatum. The Journal of neuroscience : the official journal of the Society for Neuroscience 27, 2403–2409.

Ade, K.K., Wan, Y., Chen, M., Gloss, B., and Calakos, N. (2011). An Improved BAC Transgenic Fluorescent Reporter Line for Sensitive and Specific Identification of Striatonigral Medium Spiny Neurons. Front Syst Neurosci 5, 32.

Alabi, O., Robinson, M., Fortunato, M., Kable, J.W., and Fuccillo, M.V. (2019). Disruption of Nrxn1α within excitatory forebrain circuits drives value-based dysfunction. bioRxiv.

Anderson, G.R., Aoto, J., Tabuchi, K., Foldy, C., Covy, J., Yee, A.X., Wu, D., Lee, S.J., Chen, L., Malenka, R.C., and Sudhof, T.C. (2015). beta-Neurexins Control Neural Circuits by Regulating Synaptic Endocannabinoid Signaling. Cell 162, 593–606.

Aoto, J., Martinelli, D.C., Malenka, R.C., Tabuchi, K., and Sudhof, T.C. (2013). Presynaptic neurexin-3 alternative splicing trans-synaptically controls postsynaptic AMPA receptor trafficking. Cell 154, 75–88.

Autism Genome Project, C., Szatmari, P., Paterson, A.D., Zwaigenbaum, L., Roberts, W., Brian, J., Liu, X.Q., Vincent, J.B., Skaug, J.L., Thompson, A.P., et al. (2007). Mapping autism risk loci using genetic linkage and chromosomal rearrangements. Nat Genet 39, 319–328.

Balleine, B.W., Liljeholm, M., and Ostlund, S.B. (2009). The integrative function of the basal ganglia in instrumental conditioning. Behav Brain Res 199, 43–52.

Benthall, K.N., Ong, S.L., and Bateup, H.S. (2018). Corticostriatal Transmission Is Selectively Enhanced in Striatonigral Neurons with Postnatal Loss of Tsc1. Cell Rep 23, 3197–3208.

Blundell, J., Blaiss, C.A., Etherton, M.R., Espinosa, F., Tabuchi, K., Walz, C., Bolliger, M.F., Sudhof, T.C., and Powell, C.M. (2010). Neuroligin-1 deletion results in impaired spatial memory and increased repetitive behavior. The Journal of neuroscience : the official journal of the Society for Neuroscience 30, 2115–2129.

Bucan, M., Abrahams, B.S., Wang, K., Glessner, J.T., Herman, E.I., Sonnenblick, L.I., Alvarez Retuerto, A.I., Imielinski, M., Hadley, D., Bradfield, J.P., et al. (2009). Genome-wide analyses of exonic copy number variants in a family-based study point to novel autism susceptibility genes. PLoS Genet 5, e1000536.

Budreck, E.C., Kwon, O.B., Jung, J.H., Baudouin, S., Thommen, A., Kim, H.S., Fukazawa, Y., Harada, H., Tabuchi, K., Shigemoto, R., et al. (2013). Neuroligin-1 controls synaptic abundance of NMDA-type glutamate receptors through extracellular coupling. Proceedings of the National Academy of Sciences of the United States of America 110, 725–730.

Cavaccini, A., Gritti, M., Giorgi, A., Locarno, A., Heck, N., Migliarini, S., Bertero, A., Mereu, M., Margiani, G., Trusel, M., et al. (2018). Serotonergic Signaling Controls Input-Specific Synaptic Plasticity at Striatal Circuits. Neuron 98, 801–816 e807.

Chang, J., Gilman, S.R., Chiang, A.H., Sanders, S.J., and Vitkup, D. (2015). Genotype to phenotype relationships in autism spectrum disorders. Nature neuroscience 18, 191–198.

Ching, M.S.L., Shen, Y., Tan, W.-H., Jeste, S.S., Morrow, E.M., Chen, X., Mukaddes, N.M., Yoo, S.-Y., Hanson, E., Hundley, R., et al. (2010). Deletions of NRXN1 (neurexin-1) predispose to a wide spectrum of developmental disorders. In Am J Med Genet B Neuropsychiatr Genet (Wiley Subscription Services, Inc., A Wiley Company), pp. 937–947.

Choi, K., Holly, E.N., Davatolhagh, M.F., Beier, K.T., and Fuccillo, M.V. (2019). Integrated anatomical and physiological mapping of striatal afferent projections. Eur J Neurosci 49, 623–636.

Chuhma, N., Tanaka, K.F., Hen, R., and Rayport, S. (2011). Functional connectome of the striatal medium spiny neuron. The Journal of neuroscience : the official journal of the Society for Neuroscience 31, 1183–1192.

Corbit, V.L., Manning, E.E., Gittis, A.H., and Ahmari, S.E. (2019). Strengthened Inputs from Secondary Motor Cortex to Striatum in a Mouse Model of Compulsive Behavior. The Journal of neuroscience : the official journal of the Society for Neuroscience 39, 2965–2975.

Dabell, M.P., Rosenfeld, J.A., Bader, P., Escobar, L.F., El-Khechen, D., Vallee, S.E., Dinulos, M.B., Curry, C., Fisher, J., Tervo, R., et al. (2013). Investigation of NRXN1 deletions: clinical and molecular characterization. Am J Med Genet A 161A, 717–731.

Deng, Y., Lanciego, J., Kerkerian-Le-Goff, L., Coulon, P., Salin, P., Kachidian, P., Lei, W., Del Mar, N., and Reiner, A. (2015). Differential organization of cortical inputs to striatal projection neurons of the matrix compartment in rats. Front Syst Neurosci 9, 51.

Ding, J., Peterson, J.D., and Surmeier, D.J. (2008). Corticostriatal and thalamostriatal synapses have distinctive properties. The Journal of neuroscience : the official journal of the Society for Neuroscience 28, 6483–6492.

Doig, N.M., Moss, J., and Bolam, J.P. (2010). Cortical and thalamic innervation of direct and indirect pathway medium-sized spiny neurons in mouse striatum. The Journal of neuroscience : the official journal of the Society for Neuroscience 30, 14610–14618.

Ellender, T.J., Harwood, J., Kosillo, P., Capogna, M., and Bolam, J.P. (2013). Heterogeneous properties of central lateral and parafascicular thalamic synapses in the striatum. J Physiol 591, 257–272.

Etherton, M.R., Blaiss, C.A., Powell, C.M., and Sudhof, T.C. (2009). Mouse neurexin-1alpha deletion causes correlated electrophysiological and behavioral changes consistent with cognitive impairments. Proceedings of the National Academy of Sciences of the United States of America 106, 17998–18003.

Foldy, C., Malenka, R.C., and Sudhof, T.C. (2013). Autism-associated neuroligin-3 mutations commonly disrupt tonic endocannabinoid signaling. Neuron 78, 498–509.

Fuccillo, M.V. (2016). Striatal Circuits as a Common Node for Autism Pathophysiology. Front Neurosci 10, 27.

Fuccillo, M.V., Foldy, C., Gokce, O., Rothwell, P.E., Sun, G.L., Malenka, R.C., and Sudhof, T.C. (2015). Single-Cell mRNA Profiling Reveals Cell-Type-Specific Expression of Neurexin Isoforms. Neuron 87, 326–340.

Geppert, M., Khvotchev, M., Krasnoperov, V., Goda, Y., Missler, M., Hammer, R.E., Ichtchenko, K., Petrenko, A.G., and Sudhof, T.C. (1998). Neurexin I alpha is a major alpha-latrotoxin receptor that cooperates in alpha-latrotoxin action. J Biol Chem 273, 1705–1710.

Gerfen, C.R. (1989). The neostriatal mosaic: striatal patch-matrix organization is related to cortical lamination. Science 246, 385–388.

Gerfen, C.R., Engber, T.M., Mahan, L.C., Susel, Z., Chase, T.N., Monsma, F.J., Jr., and Sibley, D.R. (1990). D1 and D2 dopamine receptor-regulated gene expression of striatonigral and striatopallidal neurons. Science 250, 1429–1432.

Giuffrida, A., Parsons, L.H., Kerr, T.M., Rodriguez de Fonseca, F., Navarro, M., and Piomelli, D. (1999). Dopamine activation of endogenous cannabinoid signaling in dorsal striatum. Nature neuroscience 2, 358–363.

Graybiel, A.M., Aosaki, T., Flaherty, A.W., and Kimura, M. (1994). The basal ganglia and adaptive motor control. Science 265, 1826–1831.

Grayton, H.M., Missler, M., Collier, D.A., and Fernandes, C. (2013). Altered social behaviours in neurexin 1alpha knockout mice resemble core symptoms in neurodevelopmental disorders. PLoS One 8, e67114.

Huang, A.Y., Yu, D., Davis, L.K., Sul, J.H., Tsetsos, F., Ramensky, V., Zelaya, I., Ramos, E.M., Osiecki, L., Chen, J.A., et al. (2017). Rare Copy Number Variants in NRXN1 and CNTN6 Increase Risk for Tourette Syndrome. Neuron 94, 1101–1111 e1107.

Hunnicutt, B.J., Jongbloets, B.C., Birdsong, W.T., Gertz, K.J., Zhong, H.N., and Mao, T.Y. (2016). A comprehensive excitatory input map of the striatum reveals novel functional organization. Elife 5.

Kattenstroth, G., Tantalaki, E., Sudhof, T.C., Gottmann, K., and Missler, M. (2004). Postsynaptic N-methyl-D-aspartate receptor function requires alpha-neurexins. Proceedings of the National Academy of Sciences of the United States of America 101, 2607–2612.

Kavalali, E.T. (2015). The mechanisms and functions of spontaneous neurotransmitter release. Nature reviews Neuroscience 16, 5–16.

Kim, H.-G., Kishikawa, S., Higgins, A.W., Seong, I.-S., Donovan, D.J., Shen, Y., Lally, E., Weiss, L.A., Najm, J., Kutsche, K., et al. (2008). Disruption of Neurexin 1 Associated with Autism Spectrum Disorder. In The American Journal of Human Genetics, pp. 199–207.

Kirov, G., Rujescu, D., Ingason, A., Collier, D.A., O’Donovan, M.C., and Owen, M.J. (2009). Neurexin 1 (NRXN1) deletions in schizophrenia. Schizophr Bull 35, 851–854.

Kreitzer, A.C., and Malenka, R.C. (2007). Endocannabinoid-mediated rescue of striatal LTD and motor deficits in Parkinson’s disease models. Nature 445, 643–647.

Kupferschmidt, D.A., Juczewski, K., Cui, G., Johnson, K.A., and Lovinger, D.M. (2017). Parallel, but Dissociable, Processing in Discrete Corticostriatal Inputs Encodes Skill Learning. Neuron 96, 476–489 e475.

Lenz, J.D., and Lobo, M.K. (2013). Optogenetic insights into striatal function and behavior. Behav Brain Res 255, 44–54.

Lin, J.Y., Lin, M.Z., Steinbach, P., and Tsien, R.Y. (2009). Characterization of engineered channelrhodopsin variants with improved properties and kinetics. Biophys J 96, 1803–1814.

Mandelbaum, G., Taranda, J., Haynes, T.M., Hochbaum, D.R., Huang, K.W., Hyun, M., Umadevi Venkataraju, K., Straub, C., Wang, W., Robertson, K., et al. (2019). Distinct Cortical-Thalamic-Striatal Circuits through the Parafascicular Nucleus. Neuron 102, 636–652 e637.

Marshall, C.R., Howrigan, D.P., Merico, D., Thiruvahindrapuram, B., Wu, W., Greer, D.S., Antaki, D., Shetty, A., Holmans, P.A., Pinto, D., et al. (2017). Contribution of copy number variants to schizophrenia from a genome-wide study of 41,321 subjects. Nat Genet 49, 27–35.

Miesenbock, G. (2011). Optogenetic control of cells and circuits. Annu Rev Cell Dev Biol 27, 731–758.

Missler, M., Zhang, W., Rohlmann, A., Kattenstroth, G., Hammer, R.E., Gottmann, K., and Sudhof, T.C. (2003). Alpha-neurexins couple Ca2+ channels to synaptic vesicle exocytosis. Nature 423, 939–948.

Packard, M.G., and Knowlton, B.J. (2002). Learning and memory functions of the Basal Ganglia. Annu Rev Neurosci 25, 563–593.

Pak, C., Danko, T., Zhang, Y., Aoto, J., Anderson, G., Maxeiner, S., Yi, F., Wernig, M., and Sudhof, T.C. (2015). Human Neuropsychiatric Disease Modeling using Conditional Deletion Reveals Synaptic Transmission Defects Caused by Heterozygous Mutations in NRXN1. Cell Stem Cell 17, 316–328.

Pan, W.X., Mao, T., and Dudman, J.T. (2010). Inputs to the dorsal striatum of the mouse reflect the parallel circuit architecture of the forebrain. Front Neuroanat 4, 147.

Reichelt, A.C., Rodgers, R.J., and Clapcote, S.J. (2012). The role of neurexins in schizophrenia and autistic spectrum disorder. Neuropharmacology 62, 1519–1526.

Rothwell, P.E., Fuccillo, M.V., Maxeiner, S., Hayton, S.J., Gokce, O., Lim, B.K., Fowler, S.C., Malenka, R.C., and Sudhof, T.C. (2014). Autism-associated neuroligin-3 mutations commonly impair striatal circuits to boost repetitive behaviors. Cell 158, 198–212.

Sara, Y., Virmani, T., Deak, F., Liu, X., and Kavalali, E.T. (2005). An isolated pool of vesicles recycles at rest and drives spontaneous neurotransmission. Neuron 45, 563–573.

Schizophrenia Working Group of the Psychiatric Genomics, C. (2014). Biological insights from 108 schizophrenia-associated genetic loci. Nature 511, 421–427.

Shen, W., Flajolet, M., Greengard, P., and Surmeier, D.J. (2008). Dichotomous dopaminergic control of striatal synaptic plasticity. Science 321, 848–851.

Smeal, R.M., Keefe, K.A., and Wilcox, K.S. (2008). Differences in excitatory transmission between thalamic and cortical afferents to single spiny efferent neurons of rat dorsal striatum. Eur J Neurosci 28, 2041–2052.

Stern, E.A., Jaeger, D., and Wilson, C.J. (1998). Membrane potential synchrony of simultaneously recorded striatal spiny neurons in vivo. Nature 394, 475–478.

Stern, E.A., Kincaid, A.E., and Wilson, C.J. (1997). Spontaneous subthreshold membrane potential fluctuations and action potential variability of rat corticostriatal and striatal neurons in vivo. J Neurophysiol 77, 1697–1715.

Sudhof, T.C. (2008). Neuroligins and neurexins link synaptic function to cognitive disease. Nature 455, 903–911.

Sudhof, T.C. (2018). Towards an Understanding of Synapse Formation. Neuron 100, 276–293.

Wall, N.R., De La Parra, M., Callaway, E.M., and Kreitzer, A.C. (2013). Differential innervation of direct- and indirect-pathway striatal projection neurons. Neuron 79, 347–360.

Willsey, A.J., Sanders, S.J., Li, M., Dong, S., Tebbenkamp, A.T., Muhle, R.A., Reilly, S.K., Lin, L., Fertuzinhos, S., Miller, J.A., et al. (2013). Coexpression networks implicate human midfetal deep cortical projection neurons in the pathogenesis of autism. Cell 155, 997–1007.

Willsey, A.J., and State, M.W. (2015). Autism spectrum disorders: from genes to neurobiology. Curr Opin Neurobiol 30, 92–99.

Wu, X., Morishita, W.K., Riley, A.M., Hale, W.D., Sudhof, T.C., and Malenka, R.C. (2019). Neuroligin-1 Signaling Controls LTP and NMDA Receptors by Distinct Molecular Pathways. Neuron 102, 621–635 e623.

Wu, Y.W., Kim, J.I., Tawfik, V.L., Lalchandani, R.R., Scherrer, G., and Ding, J.B. (2015). Input- and cell-type-specific endocannabinoid-dependent LTD in the striatum. Cell Rep 10, 75–87.

Xiong, Q., Znamenskiy, P., and Zador, A.M. (2015). Selective corticostriatal plasticity during acquisition of an auditory discrimination task. Nature 521, 348–351.

Yin, H.H., and Knowlton, B.J. (2006). The role of the basal ganglia in habit formation. Nature reviews Neuroscience 7, 464–476.

