## Supplementary figures and images for "Neurexin1α differentially regulates synaptic efficacy within striatal circuits"

### Supplemental Figures

Supp. Figure 1

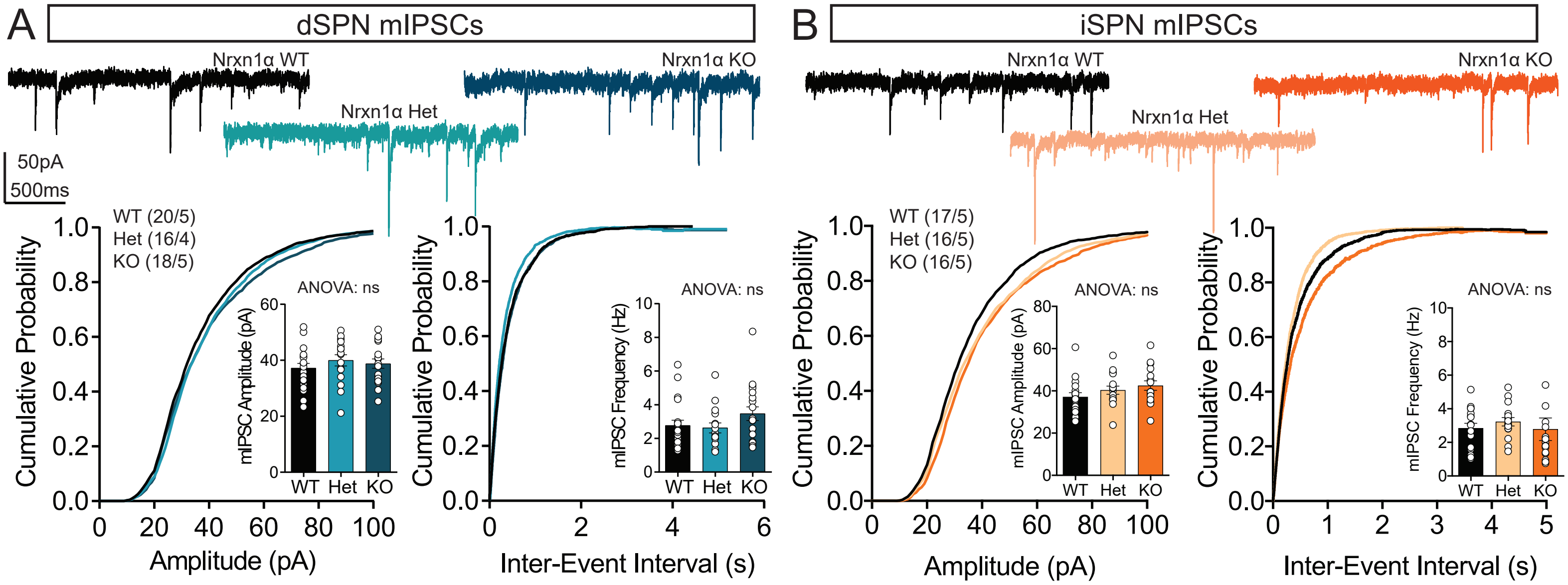

Supp. Figure 2

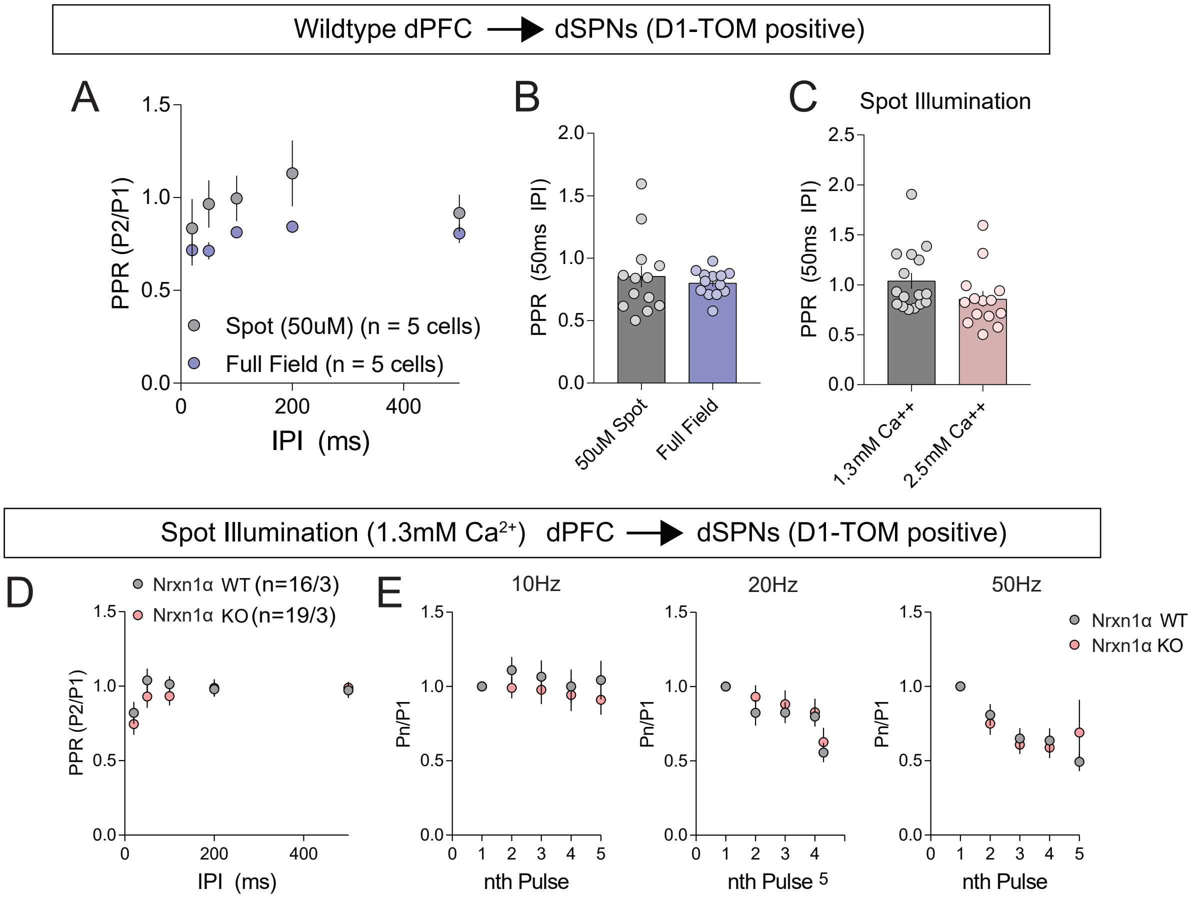

Supp. Figure 3

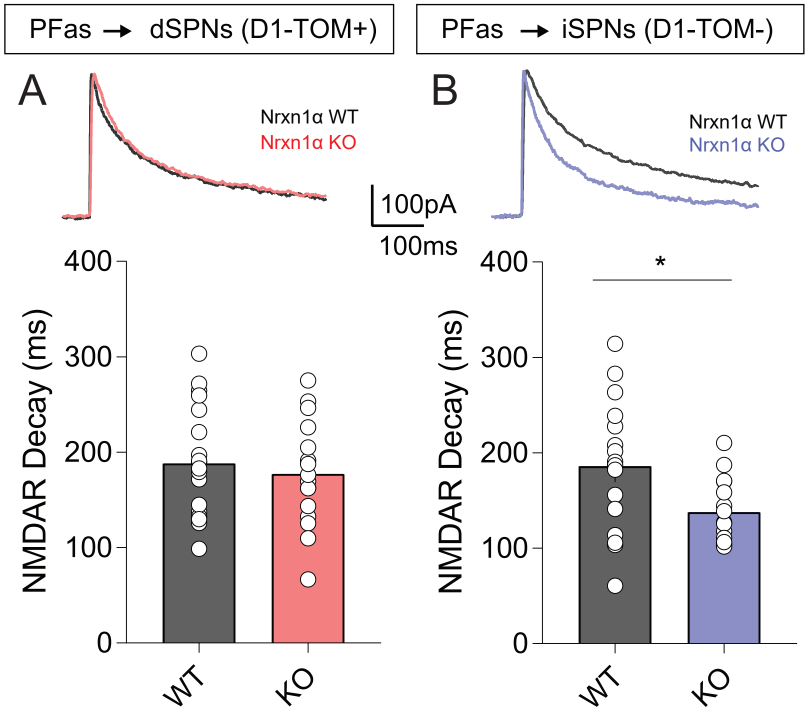

Supp. Figure 4

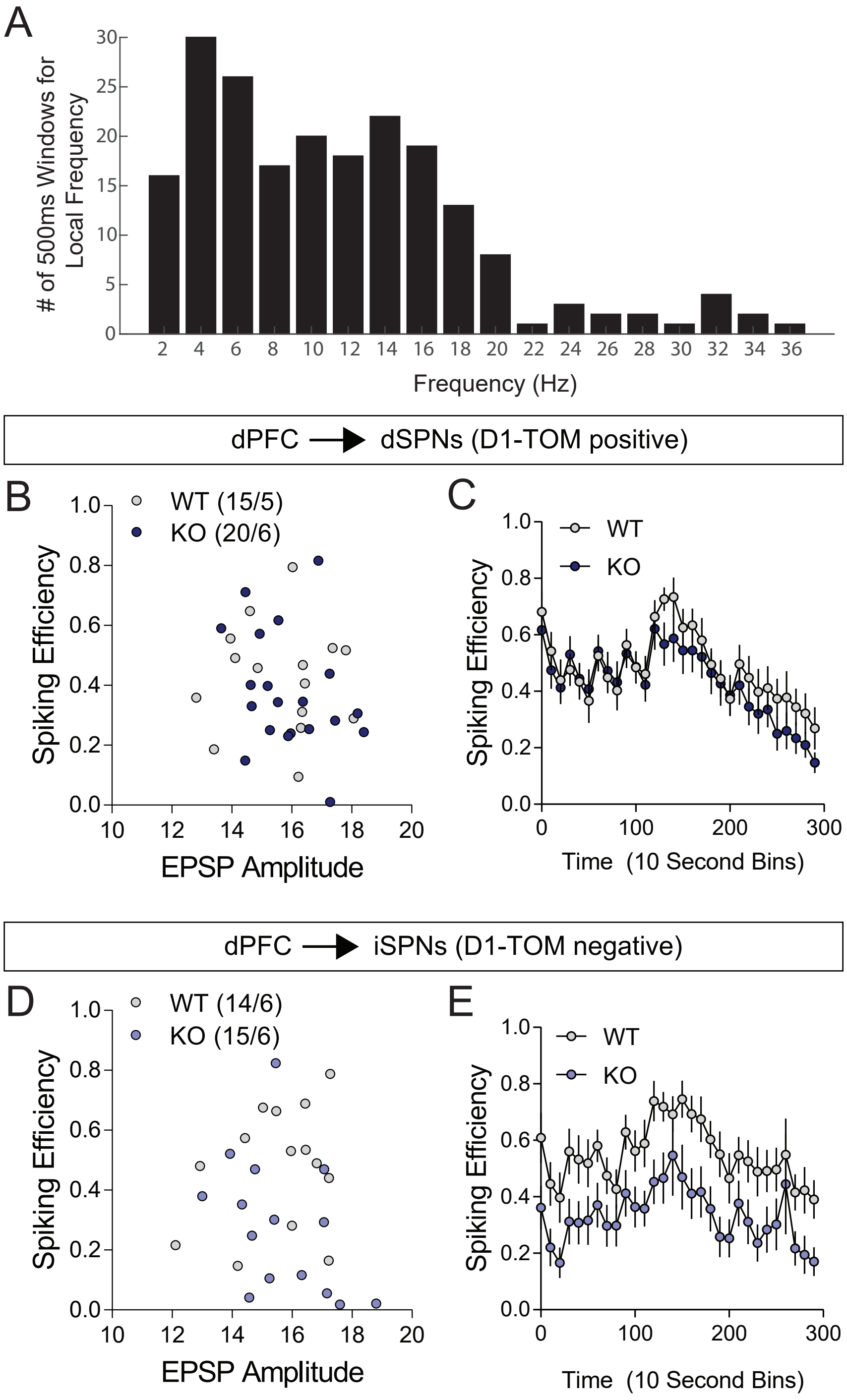
