## Supplemental Figure Legends for "Neurexin1α differentially regulates synaptic efficacy within striatal circuits"

**Supp. Figure1.** Inhibitory spontaneous synaptic transmission is unaltered in Nrxn1α mutants (related to Fig.1).

(A and B) Representative traces of mIPSCs WT, Nrxn1α Het, and Nrxn1α KO cells (top), cumulative distribution of mIPSC amplitude (lower left; inset shows average mIPSC amplitude), and cumulative distribution of inter- event intervals (lower right; inset shows average mIPSC frequency) in WT (n = 20; N = 5), Nrxn1α Het (n = 16; N = 4) or Nrxn1α KO (n = 18; N = 5) dSPNs (A) and WT (n = 17; N = 5), Nrxn1α Het (n = 16; N = 5), and Nrxn1α KO (n = 16; N = 5) iSPNs (C).

Summary data are mean ± SEM

**Supp. Figure2.** Probability of release is unchanged at dPFC-dSPN synapses regardless of external calcium levels and spot illumination measurements (related to Fig.2).

(A) Plot of paired-pulse ratio across multiple ISIs for optical stimulation of dPFC-dSPN synapses using a 50μM spot (Mightex Optical Systems) and full field illumination. Further recordings were done using an ISI of 50ms (B).

(C) Graph of paired-pulse ratio, 50ms ISI, at external calcium levels of 1.3mM and 2.5mM employing 50uM spot illumination.

(D) Plot of paired-pulse ratio across multiple ISIs for 50μM optical stimulation of dPFC-dSPN at external calcium levels of 1.3mM in Nrxn1α WT and KO.

(E) Plot of frequency trains across multiple frequencies (right; 10Hz; 20Hz; 50Hz from left to right) recorded onto dSPNs in Nrxn1α WT and KO.

Summary data are mean ± SEM

**Supp. Figure3.** Altered NMDAR decay at thalamic projections onto iSPNs in Nrxn1α KO (related to Fig5).

(A, top) Representative traces of NMDA currents measured at +40mV in the presence of 100uM picrotoxin at PFas-dSPN synapses and PFas-iSPN synapses (B, top). Traces are rescaled to same amplitude for comparison of decay time constants. Summary graph of weighted NMDAR decay values calculated from a biexponential fit for dSPNs (A) and iSPNs (B) where the fast and slow decay components are used to calculate the weighted decay time constant using the equation τ_w_= [A _f_/ A_f_ + A_s_)] x τ_f_ + [A_s_ / (A_f_ + A_s_) x τ_s._ A_f =_ fast component amplitude and A_s =_ slow component amplitude.

Summary data are mean ± SEM

**Supp. Figure4.** Initial EPSP amplitude and recording duration are similar in Nrxn1α WT and KO (related to Fig6.).

(A) Frequencies represented in an *in vivo* modeled optical stimulus pattern represented as 500ms windows and their corresponding local frequencies.

(B, D) Plot of initial EPSP (averaged 10 traces) recorded at “down-state” membrane potential of -80mV on x-axis and overall spiking efficiency on y-axis for (B) dSPNs and iSPNs (C).

(C, E) Plot of spiking efficiency across recording duration of 5 minutes consisting of 10 unique optical patterns, binned in 10-second intervals for (C) dSPNs and iSPNs recorded in Nrxn1α WT and KO.

Summary data are mean ± SEM
